# Quantitative profiling of whole-brain projectomes at single-axon resolution using deep learning and high-resolution light sheet microscopy

**DOI:** 10.1101/2025.11.14.688340

**Authors:** Ahmadreza Attarpour, Misha Raffiee, Tony Xu, Jonas Osmann, Shruti Patel, Spencer Gable-Cook, Fengqing Yu, Braedyn Au, Andrew Clappison, Mahdi Biparva, Rui Zhu, Ailey Crow, Barsin Eshaghi Gharagoz, Matthew Rozak, Isabelle Aubert, JoAnne McLaurin, Karl Deisseroth, Bojana Stefanovic, Maged Goubran

**Affiliations:** Department of Medical Biophysics, University of Toronto, Toronto, Ontario, Canada; Physical Sciences, Sunnybrook Research Institute, Toronto, Ontario, Canada; Hurvitz Brain Sciences, Sunnybrook Health Sciences Centre, Toronto, Ontario, Canada; Biological Sciences, Sunnybrook Research Institute, Toronto, Ontario, Canada; Department of Laboratory Medicine & Pathobiology, University of Toronto, Toronto, Ontario, Canada; Department of Bioengineering, Stanford University, Stanford, CA, USA; Howard Hughes Medical Institute, Stanford University, Stanford, CA, USA; Harquail centre for neuromodulation, Sunnybrook Health Sciences Centre, Toronto, Ontario, Canada

## Abstract

Revealing how individual axons contribute to a brain-wide projectome is indispensable for understanding brain function and behavior, yet remains technically challenging, with contemporary models failing to generalize. We introduce MAPL3, an end-to-end pipeline that integrates self-supervised learning with an innovative deep architecture to segment axons and reconstruct brain-wide projections in tera-voxel-scale light sheet fluorescence microscopy. Its SPECTRE network integrates local axonal features with broader information shared across neighboring 3D patches. MAPL3 generalizes across out-of-distribution datasets with different tracers, microscopes, resolutions, and protocols, outperforming state-of-the-art methods. It also provides model uncertainty estimates and improves segmentation precision on unseen data. MAPL3’s tractography module reconstructs long-range projections and quantifies passing vs. terminating streamlines. We showcase its ability to map the projectome of the orbitofrontal cortex, reconstructing streamlines that innervate established projection regions, which were missed by current pipelines. MAPL3 enables subject- and population-level quantitative profiling of whole-brain projectomes in tera-voxel-scale light sheet microscopy.

## Introduction

Without deciphering the organization of the brain-wide projectome, the complex dynamics underlying brain function and behavior will remain elusive^1–4^. At the population level, anterograde viral tracing combined with high-throughput serial two-photon tomography has enabled brain-wide, region-to-region connectivity maps in the mouse, most notably the Allen Mouse Brain Connectivity Atlas, establishing the first mesoscale reference frameworks for projectomics^3^. Complementing these population-level approaches, sparse axon reconstructions of individual projection neurons, now achievable at scales of hundreds to thousands of neurons per experiment^5–7^, have further revealed the diversity of long-range connectivity, including previously unknown neuronal subtypes and substantial heterogeneity in axonal branching, collateralization, and combinations of distal targets that are obscured by population-level measurements. Advances in tissue-clearing methods^8^, light sheet fluorescence microscopy (LSFM)^9^, and engineered adeno-associated viral vectors (AAVs) have enabled high-throughput mesoscale whole-brain connectivity imaging in intact tissue^10–13^. However, segmenting axons at single-axon resolution across tera-voxel-scale LSFM datasets remains a major technological challenge due to the immense size of these datasets, their low signal-to-noise ratio (e.g., diminished resolvability in highly arborized regions), protocol heterogeneity, artifacts, and variability in viral expression levels^1,2,14^.

State-of-the-art (SOTA) deep learning (DL) methods for LSFM analysis often fail to generalize due to biases inherent in model architectures, limited training data, and annotation variability, resulting in incomplete or spatially inaccurate reconstructions of axonal projections. Current DL-based pipelines are based on convolutional neural networks (CNNs), which struggle to extract long-range dependencies and global context, or vision transformers (ViTs), which overlook local structures and require substantial amounts of labelled data to avoid overfitting^15–17^. Hybrid CNN-transformer architectures, including TransUNet, UNETR, and Swin UNETR, combine convolutional feature extraction with transformer-based contextual modeling to improve biomedical image segmentation^15–17^. However, these architectures generally process an image or sub-volume as an independent input and do not explicitly model contextual relationships among adjacent 3D patches generated when tera-voxel LSFM datasets are partitioned for computational analysis. Although self-supervised learning (SSL) has been introduced to address the DL models’ hunger for data and has shown initial promise in microscopy^17–19^, current applications to axonal fiber mapping are limited to simple SSL frameworks and patch-level performance on small datasets^20,21^.

In biological applications of viral tracing, regional axonal projections are often quantified using coarse measures such as fluorescence intensity^22^. Similarly, current automated axonal-mapping pipelines commonly aggregate segmented axonal labeling into regional density measures at the mesoscopic scale^1,2^, potentially obscuring fine-scale and subregional projection patterns and limiting quantitative comparisons across subjects and populations. In addition, regional axonal density does not distinguish between projections that traverse a region and those whose reconstructed trajectories end within it, potentially leading regions with dense fibers of passage to be interpreted as projection endpoints. Structure tensor–based tractography of cleared-brain viral-tracing data provides complementary information by reconstructing projection streamlines^3,23^. However, these reconstructions depend on the quality and continuity of the fluorescence signal and the accuracy of local fiber-orientation estimation. Discontinuities caused by sparse labeling, signal decay, or imaging artifacts can therefore interrupt streamline propagation, producing premature or spatially inaccurate endpoints and incomplete reconstruction of distal projection pathways.

Here, we present MAPL3 (Mapping Axonal Projections in Light sheet fluorescence microscopy in 3D), an end-to-end pipeline for brain-wide axonal segmentation, streamline-based projection mapping, and quantitative population-level analysis (**Fig. 1**a). Our pipeline employs an innovative network called SPECTRE (Spatial Patch Encoding with Convolutional TransfoRmEr) (Extended Data Fig. 1 and **Fig. 1**b), with a hybrid architecture combining CNNs and ViTs that leverages patch-wise attention in addition to the standard voxel-wise attention^24^. SPECTRE extracts spatial features both between and within patches–critical for identifying global and local information, as adjacent patches share anatomical context overlooked by existing models. To mitigate DL models’ need for large datasets and scarcity of annotations, SPECTRE was pre-trained using a generative self-supervised learning (SSL)^25^ approach on nearly 22,000 unlabeled sub-volumes (256^3^ voxels, **Fig. 1**c). Model fine-tuning was done on 774 sub-volumes (Extended Data Fig. 2, 3) with annotations (almost 50,000 patches with 64^3^ voxels), using an advanced loss function designed to mitigate class imbalance, emphasize axonal edges, and improve continuity. To increase robustness further, we implemented an innovative patch-based augmentation strategy to expose the model to axons co-existing with high-intensity artifacts (hard negative patches), to focus the model on fiber morphology rather than intensity (**Fig. 1**d and Extended Data Fig. 4). We also propose pre- and post-processing algorithms (Extended Data Fig. 5), including a skeletonization algorithm, to enhance SNR of sub-volumes and fix the discontinuities of fibers due to signal decay. To estimate SPECTRE’s predictive uncertainty, we implemented Monte Carlo dropout during inference and generated voxel-wise epistemic uncertainty maps from stochastic predictions. MAPL3 further incorporates a structure tensor-based tractography module operating on voxelized segmentation maps registered to the Allen Reference Atlas. Gap-tolerant propagation enables streamlines to bridge discontinuities arising from sparse labeling or signal decay and to recover supported projections in distal regions. For each reconstructed streamline, MAPL3 records its injection-site origin, all atlas-defined regions it traverses, and its distal reconstructed endpoint. This streamline-wise annotation enables the separate quantification of passing and terminating projections. We further define the regional streamline endpoint fraction as the proportion of streamlines traversing a region that have their distal reconstructed endpoints within it, providing a traversal-normalized measure of regional endpoint concentration. By extracting regional axonal projection metrics and tract and endpoint densities, MAPL3 enables subject- and population-level statistical analysis. We validated MAPL3 across diverse test and unseen datasets (with radically different image quality and resolution) and experimental designs, demonstrating its superior generalization compared to SOTA techniques: D-LMBmap^1^ (nnU-Net-based^26^ with an axial attention module) and TrailMap^2^ (U-Net-based^27^). To highlight its utility, we applied MAPL3 to chart axonal projections originating from the orbitofrontal cortex (OFC) and contrasted it with SOTA pipelines.

**Fig 1.**
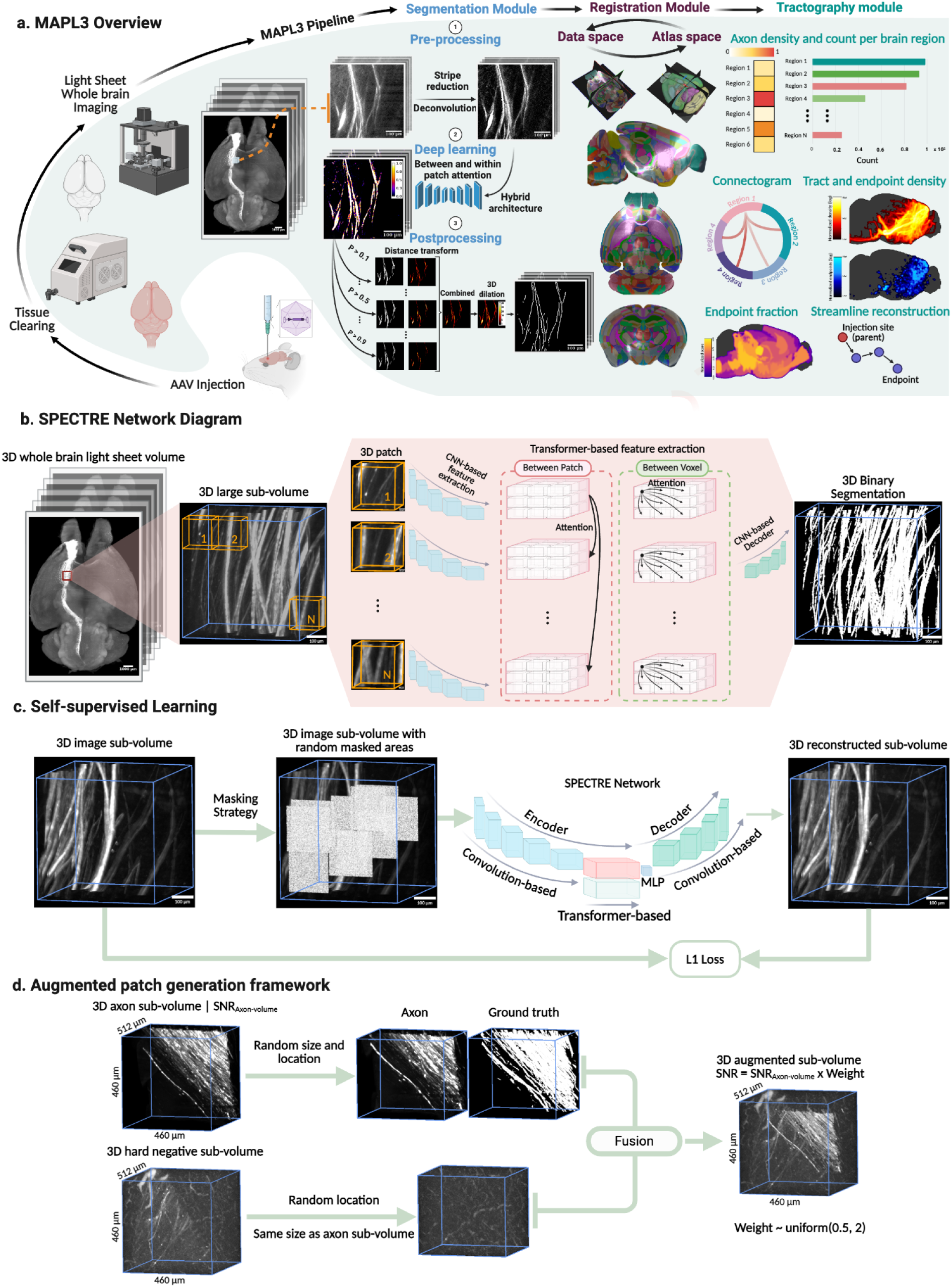
Overview of the main modules and components of MAPL3. **a.** MAPL3’s workflow. **b.** SPECTRE captures both local and global information from the large 3D sub-volume. **c.** MAPL3’s generative self-supervised learning strategy. The image on the right shows the reconstructed volume of the input on the left with randomly masked sub-volumes after five epochs. **d.** MAPL3’s patch augmentation strategy.

## Results

### MAPL3 enables brain-wide segmentation of axonal fibers at single-axon resolution in tera-voxel LSFM data

To benchmark the SPECTRE network, we first compared its performance with a SOTA vision transformer backbone architecture, UNETR16. Across all experiments on both the test and unseen datasets, SPECTRE outperformed the optimized UNETR, achieving a test DSC of 0.74 ± 0.04 compared to UNETR’s 0.69 ± 0.03 (*p* = 0.03, Mann-Whitney U test; Supplementary Table.1). The final patch-wise transformer weight in our model was 0.72, in contrast to 0.28 for the voxel-wise transformer, highlighting the importance of incorporating between-patch information in axon segmentation. Ablation studies further validated the significance of this patch-wise adaptive bias and the effectiveness of the proposed hybrid loss (Supplementary Table 2).

Pretraining benefited our MAPL3 pipeline over random initialization, improving the test DSC to 0.77±0.02 (4.1% increase) and yielding a 9.4% increase in precision on the unseen dataset (*p* = 0. 04 Mann-Whitney U test; Supplementary Table 3), underscoring the enhanced generalizability conferred by SSL. On both the test and unseen datasets, MAPL3 (without post-processing) outperformed SOTA methods across all evaluation metrics (**Fig. 2**a), highlighting its superior generalizability in new experimental protocols and in regions containing both sparse and dense fibers (e.g., MAPL3 achieved a DSC improvement of 0.30 vs. D-LMBmap, *p* = 0. 0001; and 0.50 vs. TrailMap, *p* = 0. 0001; Mann-Whitney U test). While MAPL3 demonstrated consistent performance across in- and out-of-distribution datasets, D-LMBmap outperformed TrailMap on in-distribution data but underperformed on out-of-distribution data (**Fig. 2**b). We further fine-tuned TrailMap and D-LMBmap using the same annotated training data used for MAPL3 and evaluated all three models on the test and unseen datasets. MAPL3 retained significantly higher DSC and lower 95th-percentile Hausdorff distance than both fine-tuned benchmarks on both datasets, achieving an average relative DSC improvement of more than 25% on the unseen dataset (Fig. 2b). Fine-tuning increased TrailMap recall but resulted in substantially lower precision, whereas D-LMBmap achieved precision comparable to MAPL3 but exhibited lower recall on both datasets. Moreover, MAPL3 demonstrated strong generalization to LSFM datasets acquired using different AAV serotypes and fluorophores, and imaging setups, such as TrailMap benchmarking dataset and D-LMBmap’s DCN Stanford dataset (Extended Data Fig. 6), likely due to the diverse representations learned during SSL and data augmentations.

**Fig 2.**
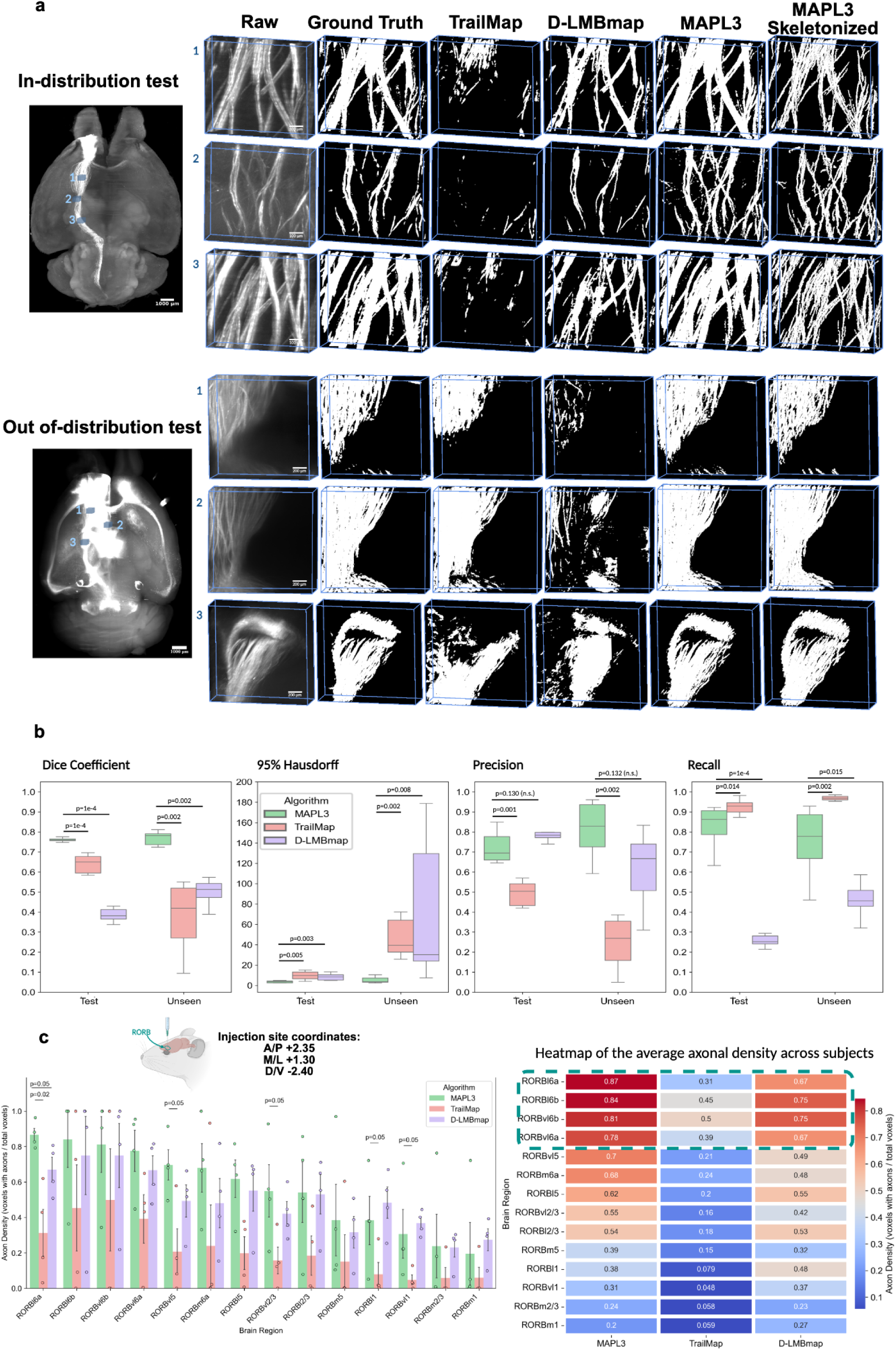
Performance of MAPL3 in brain-wide segmentation of axonal fibers compared to SOTA methods. Qualitative (**a)** and quantitative (**b)** comparison of segmentation accuracy between MAPL3, TrailMap, and D-LMBmap on in- and out-of-distribution test datasets. Box plots: box limits, upper and lower quartiles; center line, median; whiskers, 1.5× interquartile range; points, outliers. N=2112 unique patches ( 64^3^*voxels* ≃ 0. 11^3^*mm*^3^) and N=384 unique patches of 64^3^*voxels* ≃ 0. 32^3^*mm*^3^ for test and unseen datasets, respectively. Mann-Whitney U test (two-sided). **c.** *eYFP+* cell density per label plotted for each subdivision in the right OFC (left panel), averaged across subjects, and visualized as a heatmap, ordered from high to low density (right panel). The green dashed box highlights the top subdivisions in terms of axonal density.

To estimate model uncertainty during axonal segmentation, we applied Monte Carlo dropout to the unseen datasets by retaining dropout layers during inference and performing 50 stochastic forward passes for each input sub-volume (Supplementary Data Fig. 2). The mean probability across predictions was used to generate the ensemble segmentation, whereas the voxel-wise standard deviation represented epistemic uncertainty in the model predictions. As expected, uncertainty was concentrated predominantly around axonal boundaries and regions containing weak or ambiguous signal, while consistently detected axons exhibited lower variability across stochastic predictions (Supplementary Data Fig. 2). Compared with deterministic MAPL3 inference, MC-dropout ensemble prediction increased precision across the unseen samples, corresponding to an average relative precision gain of approximately 8.1% over the MAPL3 baseline. These findings demonstrate that repeated stochastic inference provides spatially resolved estimates of model uncertainty while reducing false-positive segmentation in images acquired under previously unseen experimental conditions.

### Quantitative mapping of brain-wide OFC projection patterns

To validate MAPL3’s ability to identify individual fibers and reconstruct their putative brain-wide projections, we applied our pipeline to map the OFC circuit in unseen LSFM datasets with low and high image resolutions (scanned on different microscopes), where AAV8-CaMKIIa-eYFP-NRN was unilaterally injected into the right OFC (or ORB in Allen reference atlas ‘ARA’). Group-wise heatmaps and regional bar plots of segmented axonal density within the injection site and its subdivisions revealed MAPL3’s ability to identify consistent labeling across subjects, with the lateral (ORBl6a-b) and ventrolateral (ORBvl6a-b) areas showing the strongest signals–consistent with the targeted stereotaxic coordinates (**Fig. 2**c). In addition to demonstrating the lowest within-region standard deviation, MAPL3 captured a 4-fold variation in axonal density across OFC subdivisions (0.20-0.87), reflecting biological heterogeneity in viral uptake and projection strength; while, TrailMap and D-LMBmap produced compressed dynamic ranges (0.04-0.50 and 0.23-0.75, respectively), indicating reduced sensitivity to regional differences (**Fig. 2**c).

Transforming the segmented axonal projectomes and raw fluorescence images into ARA space (**Fig. 3**a-b) demonstrated that MAPL3 (without fine-tuning) maintained fiber continuity across long-range projections from the right orbital area (RORB) to the Caudoputamen and then to the Midbrain region, while avoiding false positives (**Fig. 3**b and Supplementary Data Fig. 4). MAPL3 accurately delineated fibers within the corpus callosum and contralateral OFC that were missed or incompletely captured by TrailMap and D-LMBmap (Extended Data Fig. 7, 8). In high-intensity regions that receive the most innervation from OFC, D-LMBmap tended to produce widespread false positives, whereas TrailMap failed to detect axonal fibers, likely reflecting differences in training data and loss formulation between the two methods (Extended Data Fig. 9). In challenging, fiber-sparse regions, such as the mediodorsal nucleus of thalamus (MD) and periaqueductal gray (PAG)^22^, MAPL3 accurately reconstructed individual axonal fibers, whereas TrailMap exhibited low recall and D-LMBmap produced widespread over-segmentation (**Fig. 3**c and Supplementary Data Fig. 5).

**Fig 3.**
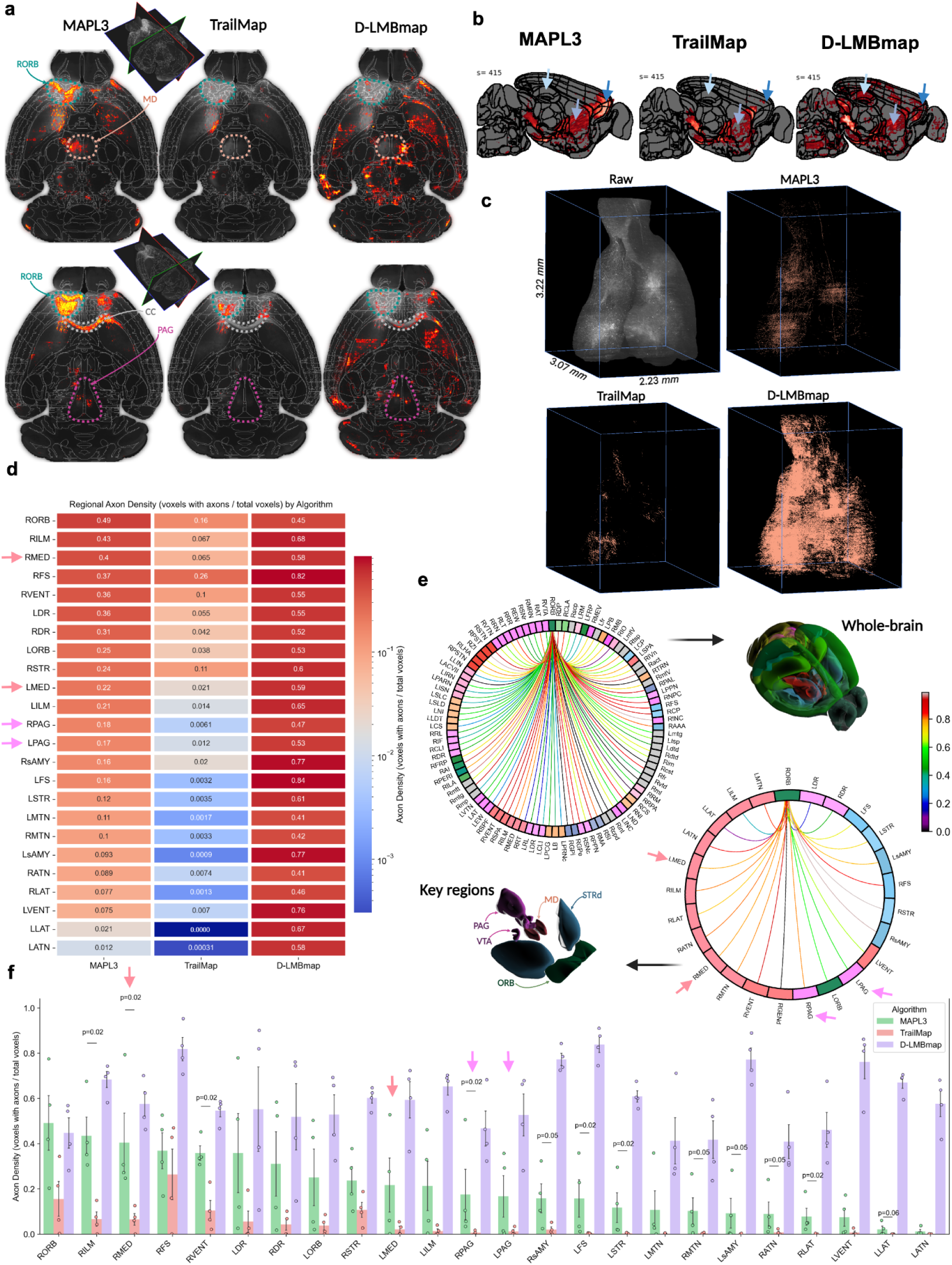
Brain-wide projection mapping and regional density analysis of OFC circuitry. Axial (**a**) and sagittal (**b**) slices showing ARA-aligned segmentation maps overlaid on labels, visualizing outputs from MAPL3, TrailMap, and D-LMBmap alongside the warped raw images. In **a**, key regions are highlighted, including RORB, MD, PAG, and corpus callosum (CC). In **b**, arrows highlight the continuity and axonal collateral map detected by MAPL3. **c.** PAG was identified in native coordinates and cropped from the raw image and segmentation outputs of each method to assess regional accuracy. Segmented *eYFP+* axonal densities per label (N=4/group) shown as bar plots (**d;** two-sided Mann–Whitney U tests) and a regional heatmap (**f**) for key regions at depth 6. Arrows highlight differential detection in the MD and PAG regions. **e.** Regional connectivity is visualized as circular plots.

Quantitative analysis demonstrated that MAPL3 uncovered more comprehensive and anatomically precise regional projection patterns, capturing dense projections to key validated target regions, such as the MD and PAG, as well as contralateral cortical areas (Supplementary Data Fig. 6). The dense axonal labeling detected in the MD aligns with previous studies reporting the role of mOFC→MD connectivity in alcohol-activated neuronal ensembles, highlighting MAPL3’s ability to resolve biologically meaningful projections^22^. MAPL3’s quantification of axonal density at an ARA depth 6 (atlas hierarchy) revealed a smooth decline from the injection site to distal targets, including the right and left mediodorsal thalamic nuclei (RMED and LMED), and striatum, which mirrors the expected spatial distribution of OFC axonal projections ^28–30^ (**Fig.3** d–f). Extending this analysis at the whole-brain level (all ARA subdivisions), MAPL3 identified high projection densities to the bilateral dorsal raphe nucleus (DR) and ventral tegmental area (VTA), consistent with prior tracing studies showing that OFC projects to dopaminergic neurons in these regions and midbrain monoaminergic nuclei^28,31,32^ (**Fig. 3**e and Supplementary Data Figs. 7, 8).

Distinct projections from the prefrontal cortex to PAG mediate behaviorally specific functions, including analgesic and anxiolytic effects of the dorsal medial prefrontal cortex to the ventrolateral PAG pathway^33^. To perform quantitative population-level analysis, we normalized regional axonal density to whole-brain axonal density. Investigating OFC→MD projections at the sub-regional level, MAPL3 delineated localized patterns of sub-regional axonal labeling and preserved anatomically consistent a rostrocaudal–ventrodorsal gradients of axonal innervation across PAG subdivisions^34,35^; with high densities in subregions such as the Interstitial nucleus of Cajal and nucleus of Darkschewitsch (ND) that closely reflected raw fluorescence patterns and were consistent with previous studies^34,35^ (Extended Data Fig.10). In contrast, D-LMBmap produced regional projection-density profiles due to diffuse false positives, and TrailMap often failed to detect projections in individual brains, ultimately resulting in flawed connectivity (Extended Data Fig. 10a). MAPL3 further enabled reproducible mapping of these anatomically localized projection patterns with high cross-subject consistency (Extended Data Fig. 10b and Supplementary Data Fig. 9).

### AI-based reconstruction of OFC projections

Regional axonal density quantifies the distribution of (segmented) axonal labeling but does not distinguish projections traversing a region from those whose reconstructed trajectories end within it. To operationally separate these projection features, we extended MAPL3 with a streamline-based tractography module operating on voxelized segmentation maps (registered to ARA space; **Fig. 4**a). Structure tensor analysis (STA) was used to estimate local fiber coherence and principal orientation from the *segmented axonal signal*. Streamlines were then propagated from the right orbital area (RORB) injection site using constraints on step size, gap tolerance, turning angle, curvature, and total streamline length (Fig. 4a).

**Fig 4.**
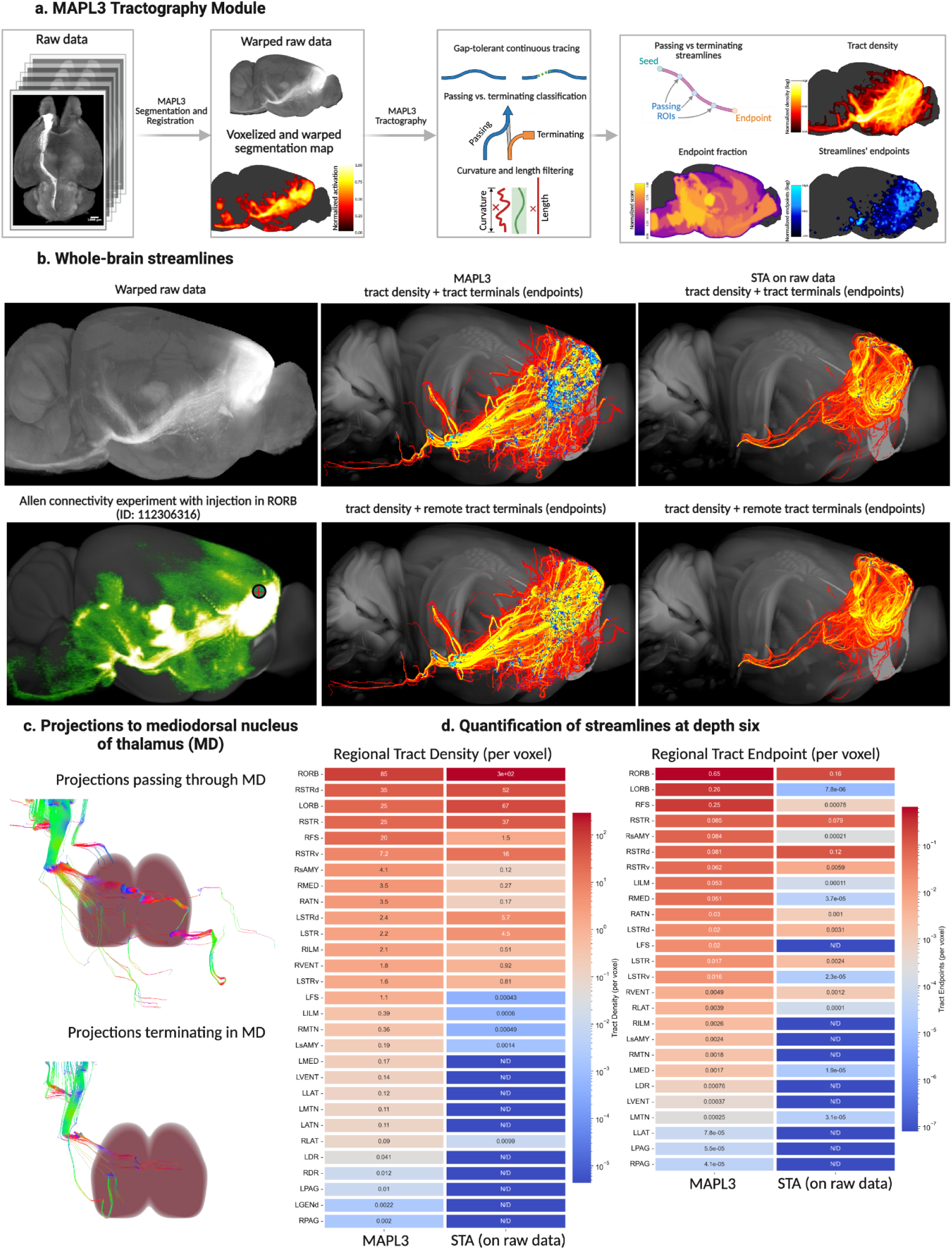
MAPL3 tractography reconstructs whole-brain axonal projections and distinguishes terminal fields from fibres of passage. **(a)** Overview of the MAPL3 tractography module. Raw light-sheet volumes are segmented and registered to the Allen atlas at 25 µm isotropic resolution, yielding a voxelized and warped segmentation map from which a structure tensor field is computed. Streamlines are then propagated by gap-tolerant continuous tracing, and are refined by curvature and length filtering. The module outputs tract density, streamline endpoint density, and the regional streamline endpoint fraction (EF), the proportion of streamlines traversing a region that terminate within it, which separates regions that receive terminal projections from those crossed by fibres en route to more distal targets. **(b)** Whole-brain reconstruction from RORB. Left, warped raw signal (top) and a comparable Allen Mouse Brain Connectivity experiment (ID 112306316; bottom), with the injection site circled. Center and right, tract density overlaid with streamline endpoints for MAPL3 and for structure tensor analysis (STA) applied to the raw data, shown with all endpoints (top) and with endpoints distal to the injection site only (bottom). MAPL3 recovers a more extensive and continuous projection field, including long-range descending pathways that STA does not reconstruct. **(c)** Streamlines intersecting the mediodorsal nucleus of the thalamus (MD), separated into those passing through MD and routing elsewhere (top) and those terminating within it (bottom), illustrating the distinction that tract density alone cannot resolve. **(d)** Regional quantification at ontology depth six, comparing MAPL3 with STA for tract density per voxel (left) and streamline endpoints per voxel (right). Values are log-scaled; N/D denotes regions in which the algorithm detected no streamlines. MAPL3 detects projections in numerous regions that STA does not resolve, including the medial group of the dorsal thalamus (RMED, LMED), periaqueductal gray (RPAG, LPAG), and dorsal raphe (RDR, LDR).

We applied the tractography module to representative brains and compared the resulting tractography-derived pathways with those generated by applying the same STA-based procedure to the downsampled LSFM fluorescence signal registered to ARA space, an approach previously used to reconstruct tracts from cleared-brain viral-tracing data. Both analyses used the same injection-site mask, streamline seeds, and tracking parameters. For each reconstructed streamline, MAPL3 recorded the injection-site origin, all atlas-defined regions it traversed, and its distal atlas-defined endpoint. Operating on the voxelized MAPL3 segmentation maps, together with the implemented gap-tolerance strategy, enabled the reconstruction of streamlines extending to established OFC projection regions distal to the injection site such as the PAG, MD, and VTA that were missed when tractography was performed directly on the downsampled fluorescence signal (**Fig. 4**b and Supplementary Data Fig. 10). Whole-brain tractograms further demonstrated the broader spatial extent of MAPL3 segmentation-based trajectories compared with trajectories reconstructed directly from the downsampled fluorescence signal (Supplementary Data Fig. 10). To visualize region-specific projections and the effect of passing vs. terminating tracking, we isolated streamlines that either traversed the MD (**Fig. 4**c) and VTA (Supplementary Data Fig. 11) or had their distal reconstructed endpoints within it. We subsequently quantified the tract density index and streamline endpoint density within each ARA label and compared the regional profiles obtained from MAPL3 segmentation-based and fluorescence-signal-based tractography at ARA depth 6 (**Fig. 4**d) and across all brain regions (Supplementary Data Fig. 12). This analysis revealed more comprehensive regional projection profiles, including trajectories extending to the DR, and provided streamline-wise regional measurements that are not generated by the standard TrailMap or D-LMBmap workflows. These tractography-derived pathways were spatially concordant with MAPL3-segmented axonal labeling, the corresponding fluorescence signal, and projection patterns reported for comparable experiments in the Allen Mouse Brain Connectivity Atlas (Fig. 4b and Extended Data Fig. 10).

Based on MAPL3’s tractography we computed an endpoint density metric, defined as the number of distal reconstructed endpoints per voxel within each atlas-defined region, and mapped its whole-brain distribution (Extended Data Fig. 10b,e, and f). High endpoint densities were observed outside the injection site in established OFC projection regions, including the medial group of the dorsal thalamus (such as MD, and Medial group of the dorsal thalamus, SMT), and VTA (Extended Data Fig. 10e). Endpoint-density analysis also resolved differences among atlas-defined subdivisions of the injection site, highlighting biological heterogeneity in projection strength (Extended Data Fig. 10f and Fig. 2c). In addition to endpoint density, we computed streamline endpoint fraction (EF) as the number of streamlines with endpoints within a region divided by the number of streamlines traversing that region (Extended Data Fig. 10a,c,d,g-i). EF showed no detectable association with distance from the injection location (Spearman’s ρ = +0.08, 95% CI −0.02 to +0.18), whereas tract density and endpoint density per voxel both covaried strongly with proximity (ρ = −0.56 and −0.54, respectively; both P < 0.0001). EF was only weakly associated with tract density (ρ = −0.16, 95% CI −0.26 to −0.06), indicating that EF and tract density rank regions differently.

As a proof-of-concept anatomical evaluation of the utility of these metrics, we compared their ability to differentiate a curated reference set of white matter fiber tracts vs. established gray matter OFC projection regions. EF differentiated the two reference sets with an area under the receiver operating characteristic curve of 0.849 at a permissive sampling threshold of at least 50 traversing streamlines (P = 1.5 × 10⁻⁴, two-sided Mann–Whitney U test; Extended Data Fig. 10g). In contrast, the areas under the curve were 0.280 for tract density and 0.475 for endpoint density. Mean EF was three- to fivefold higher in the gray matter regions than in the white matter fiber tracts reference set (Extended Data Fig. 10h). The complementary information provided by EF was also evident among regions with high traversal counts. The right preparasubthalamic nucleus (RPST), right parasubthalamic nucleus (RPSTN), and right subthalamic nucleus (RSTN) contained 21,599, 51,253, and 47,246 traversing streamlines, respectively, but few or no distal reconstructed endpoints (EF = 0.00, 0.01, and 0.04, respectively; Extended Data Fig. 10i). These regions are gray matter nuclei, demonstrating that regions dominated by traversing streamlines are not restricted to major white matter pathways. Collectively, these findings demonstrate that EF provides information complementary to regional tract and endpoint density by identifying whether a region is predominantly traversed by reconstructed streamlines or enriched in distal reconstructed endpoints. Nevertheless, EF quantifies the regional concentration of computationally reconstructed distal endpoints and does not directly identify biological axon terminals, presynaptic specializations, or synaptic connectivity.

## Discussion

This work introduces MAPL3, an end-to-end pipeline for native-resolution axon segmentation, atlas-based quantification, and streamline-wise reconstruction of brain-wide mesoscopic projectomes in tera-voxel-scale LSFM data. MAPL3 achieved precise, generalizable segmentation across heterogeneous high-resolution LSFM datasets acquired using different tracers, microscopes, spatial resolutions, clearing methods, and imaging protocols. This performance is supported by generative self-supervised pretraining, SPECTRE’s parallel voxel-wise and patch-wise transformer branches, a hybrid loss function, and hard-negative patch augmentation. Comparisons with fine-tuned TrailMap and D-LMBmap models on the same training data demonstrated that MAPL3’s performance gains persisted across in- and out-of-distribution datasets, while ablation experiments further established the contributions of SPECTRE’s architectural and task-specific components. Monte Carlo dropout additionally provided voxel-wise uncertainty estimates and improved precision on unseen datasets through stochastic ensemble inference. Together, these features establish MAPL3 as a modular, accessible pipeline for native-resolution, brain-wide projectome mapping and comparative connectivity analysis across subjects and experimental cohorts.

Application of MAPL3 to the RORB projectome recovered anatomically localized projection patterns across cortical, striatal, thalamic, hypothalamic, and midbrain regions. These regional distributions were consistent with population-level tracing and single-neuron reconstruction studies demonstrating extensive and heterogeneous OFC projections throughout the mouse brain^3,36,37^. Whole-brain reconstructions have further shown that individual OFC projection neurons exhibit divergent, layer-dependent projections, emphasizing the fine-scale organization obscured when axonal labeling is summarized solely using regional intensity or density measurements^36^. (Extended Data Fig. 9). MAPL3 identified dense axonal labeling in established OFC projection regions, including nucleus accumbens, mediodorsal and submedial thalamic nuclei, and midbrain structures. Projections toward the mediodorsal and submedial thalamic nuclei were consistent with established reciprocal relationships between the orbital cortex and medial thalamus^38,39^, whereas the projection toward the PAG agreed with tracing and functional evidence for a direct vlOFC-to-vlPAG pathway^40^. The broader cortical and subcortical distribution was also concordant with comparable experiments in the Allen Mouse Brain Connectivity Atlas^3^.

Regional axonal density cannot alone determine whether labeled projections traverse or end within a region. Previous STA tractography of cleared-brain viral-tracing data operated directly on downsampled and registered fluorescence images and showed that input-signal quality and continuity, fiber-orientation estimation, and stopping criteria influence the resulting reconstructed projections and endpoint locations^23^. By performing tractography on MAPL3-segmented axonal signal, our pipeline reduces the contribution of nonspecific fluorescence and high-intensity artifacts to orientation estimation. Optimized gap-tolerant propagation for whole-brain LSFM further enables streamlines to bridge short discontinuities caused by sparse labeling or signal decay, while constraints on step size, turning angle, curvature, and trajectory length limit anatomically implausible trajectories. Compared with the same STA procedure applied to the downsampled LSFM fluorescence signal, MAPL3 segmentation-based tractography reconstructed more complete distal pathways and recovered trajectories extending to known OFC-associated regions, including the dorsal raphe nucleus, that were missed using the fluorescence-based input and exclusion of gap tolerance.

The streamline endpoint fraction (EF) metric provided complementary information to tract and endpoint density by distinguishing major white matter fiber tracts from established gray matter OFC projection regions and revealing that several densely traversed gray matter regions were not enriched in distal reconstructed endpoints. Streamline-wise annotation therefore extends projectome analysis beyond regional density measurements by enabling filtering of reconstructed pathways by origin, traversed regions, and distal endpoints. However, EF quantifies the regional concentration of distal reconstructed streamline endpoints and does not directly identify biological axon terminals, presynaptic specializations, or synaptic connectivity.

While MAPL3 performs native-resolution segmentation and registration of axons in tera-voxel LSFM datasets, our expert-corrected ground truth was binary semantic labels and did not assign unique identities to individual axons. The current evaluation therefore does not quantify instance-level false merges or splits, or branch-point accuracy throughout the brain. Moreover, voxelization and structure tensor estimation may combine orientation information from multiple adjacent axons, particularly within dense or crossing bundles. The resulting streamlines should therefore be interpreted as putative projections rather than one-to-one reconstructions of individual biological axons, and their endpoints should not be considered direct evidence of axonal terminals or synaptic connectivity. Tractography also remains dependent on segmentation continuity, gap-tolerance parameters, orientation estimation, and streamline stopping criteria. Future studies incorporating synaptic markers, manual whole-axon reconstructions, or higher-resolution imaging could provide instance-level and synaptic ground truth.

In summary, MAPL3 provides an end-to-end framework for native-resolution segmentation, anatomical mapping, and quantitative analysis of axonal projectomes in tera-voxel-scale LSFM data. Integrating SPECTRE with self-supervised pretraining, predictive uncertainty estimation, atlas registration, and streamline-based tractography provides a unified pipeline to investigate regional axonal density, streamline traversal, endpoint distributions, and inter-subject variability. Although developed and validated for fluorescently labeled axonal projections, the methodological advances introduced here, particularly SPECTRE’s adaptive integration of within-patch morphology and contextual information across adjacent 3D patches, are transferable to other large-scale volumetric imaging domains following task-specific fine-tuning. MAPL3’s modular design also facilitates integration of alternative segmentation models, anatomical atlases, and downstream analytical approaches. MAPL3 provides an accessible, expandable framework for investigating brain-wide axonal organization across experimental subjects and populations and can support broader applications in projectomics, large-scale volumetric imaging, and connectivity mapping.

## Acknowledgments

This work was supported by funding from Canadian Institute of Health Research (CIHR) grants PJT178059, PJT376309, and PJT156179, National Institutes of Health (NIH) grant R01AG084681-01, Natural Sciences and Engineering Research Council (NSERC) Discovery grant RGPIN-2021-03728, and the Alzheimer’s Society young investigator grant #23-05. A.A. was funded by the Department of Medical Biophysics, University of Toronto’s Excellence Award. B.S. is funded by the Canada Research Chairs program award #CRC-2018-00042. M.G. is supported by the Canada Research Chairs program award #CRC-2021-00374. We are grateful to the Digital Research Alliance of Canada (alliancecan.ca) for their allocation of computing resources used in parts of this research. We are grateful for support from the Harquail Centre for Neuromodulation.

## Author contributions

AA, BS, and MG conceived and led the development of MAPL3. AA developed and evaluated all the deep learning models, conducted all the data processing and feature extraction analysis, and created the figures, with input from MG and BS. TX, FY, MB, and RZ contributed to DL model development, optimization, and data processing. MRa, ACr, and BG conducted the experiments, performed data acquisition, and contributed to the interpretation of the results. JO, AA, and MG developed and implemented the MAPL3 Docker and integration with the MIRACL platform. SP, BA, and MRo assisted with ground truth data generation and contributed to the interpretation of results. ACl contributed to DL model evaluation. IA, JM, and KD contributed to the experimental design and the interpretation of results. AA, BS, and MG wrote the manuscript with input from all the co-authors. MG and BS received funding and supervised all aspects of this work.

## Competing interests

The authors declare no competing interests.

## Code availability

The MAPL3 pipeline, along with the SPECTRE network, will be publicly available (upon acceptance) as an end-to-end module within the open-source MIRACL platform (CC BY-NC-ND 4.0 license) at https://miracl.readthedocs.io/, together with documentation, tutorials, example data, graphical user interfaces, and visualization tools. MAPL3 is implemented in a modular fashion, with well-documented core functions that can be executed either as command-line tools or through MIRACL GUIs. Available modules include segmentation (with user-friendly fine-tuning scripts), voxelization, registration, and statistical analysis.

## Data availability

Different versions of the pretrained deep learning models are provided with the pipeline. A subset of datasets generated and analyzed in this study (voxelized and warped segmentation maps) is available at www.miracl.readthedocs.io under the MAPL3 workflow tutorial as example data. All required resources, including the Allen Reference Atlas, anatomical labels, and datasets, are packaged in the MIRACL containers, along with documentation and tutorials.

## Methods

### Animals

All animal procedures met the guidelines of the National Institutes of Health and were approved by Stanford University’s Institutional Animal Care and Use Committee. Adult male C57BL/6J mice (Jackson Laboratory, strain #000664) at 8-12 weeks of age were used.

### Stereotaxic Surgery

Mice were anesthetized with 1-3% isoflurane and placed in a small animal stereotaxic apparatus (Kopf Instruments). Body temperature was maintained at 37°C using a closed-loop temperature control system combining a flexible rectal probe for measurement with an external heating pad (Homeothermic Monitoring System, Harvard Apparatus). For anterograde labeling of axonal projections, AAV8-CaMKIIa-eYFP-NRN (1 x 10^12^ vg/mL, Stanford Neuroscience Gene Vector and Virus Core) was unilaterally injected in the right orbitofrontal cortex at A/P +2.35, M/L +1.30, and D/V -2.40 at a rate of 100 nL/min.

### Datasets

Our pre-training and training data consisted of 11 animals, with five used for the pre-training stage and six animals used for training. For training, four animals were allocated to the training set, with one animal assigned to each of the validation and test sets. We further employed four animals for our application dataset. All of these animals were cleared with SHIELD^41,42^ and imaged using a SmartSPIM microscope (see below). For unseen evaluation, we used a whole-brain dataset from one animal that was cleared with SHIELD and imaged with a LaVision microscope (see below).

### Tissue clearing and Immunolabeling

Mice were injected with sodium pentobarbital (200 mg/kg) and transcardially perfused with ice-cold 1X phosphate-buffered saline (PBS) and 4% paraformaldehyde (PFA) in PBS. Whole-brain samples were post-fixed in SHIELD GE38 epoxy in 4% PFA (LifeCanvas Technologies) and processed with commercial whole-brain clearing methods (SHIELD Kit, LifeCanvas). Samples were then electrophoretically cleared for 3 days using a SmartBatch+ (LifeCanvas) and washed four times for 12 hrs in 1X PBS with 0.3% Triton (PBST) at 37°C.

### Light sheet fluorescence microscopy (LSFM) imaging

#### SmartSPIM Acquisition

Following clearing, whole-brain samples were index-matched with gentle shaking overnight in EasyIndex solution (RI = 1.52) at RT. Samples were subsequently mounted individually in 1% EasyIndex agarose gel, acclimated overnight to EasyIndex imaging solution in the imaging chamber, and imaged on a LifeCanvas SmartSPIM with 1.8um XY resolution and 3µm Z resolution (3.6X objective). Post-processing of whole-brain images was performed to remove striping artifacts (PyStripe), tiling artifacts by modeling illumination as a Gaussian and defining parameters of sigma and amplitude to normalize light spread across each tile (LifeCanvas), and stitching acquired axial view tiles per Z plane (Terastitcher^43^).

#### LaVision Acquisition

Whole-brain cleared samples were acclimated overnight with gentle shaking at room temperature in refractive index-matching solution, sPROTOS^44^. sPROTOS was prepared by combining 40g N-methyl-D-glucamine, 55g iohexol, and 50g diatrizoic acid in 100 mL UltraPure DNase/RNase-Free Distilled Water, and adjusting RI to 1.47. Samples were mounted directly on a custom, 3D-printed sample holder with adhesive and acclimated overnight in the imaging chamber. Samples were imaged in sPROTOS on a LaVision Ultramicroscope II with a 10x objective and 0.63x zoom macro lens at 5μm isotropic resolution.

### MAPL3 algorithm and workflow

MAPL3 enables a robust analysis of single axonal fibers and their whole-brain projections in tissue-cleared LSFM datasets (**Fig. 1**a). Whole-brain images are first divided into smaller 256³ sub-volumes within the MAPL3 segmentation module; in preprocessing, ‘stripe’ and ‘shadow’ artifacts commonly associated with LSFM datasets are removed and SNR enhanced in each image patch by smoothing out the background (Extended Data Fig. 5a). Axonal fibers are segmented from the preprocessed volumes using SPECTRE network (**Fig. 1**b and Extended Data Fig. 1). Local features are extracted from each 64³ sub-volume using a series of five optimized convolution-based encoder blocks and then processed by two parallelized transformer blocks with a shortcut connection: a voxel-wise transformer, which models fine-grained relationships within each sub-volume, and a patch-wise transformer, which captures attention (shared information) across multiple sub-volumes. The extracted features are then concatenated, weighted, and input into a convolutional-based decoder to restore spatial resolution. Since adjacent image patches in the LSFM dataset share common features, we designed a globally aware bias term using the patch-wise transformer. The patterns and information obtained from the patch-wise transformer is incorporated into the output of the decoder (Extended Data Fig. 1). The resulting segmentation probability maps are thinned and filtered based on morphology to minimize discontinuities (Extended Data Fig. 5b). Using our MIRACL platform, whole-brain data are then registered to the Allen Mouse Brain Reference Atlas (ARA), through linear and nonlinear transformations (**Fig. 1**a). Deformation fields derived from this registration, along with atlas labels and binarized segmentation maps, are processed by the MAPL3 feature extraction module to produce quantitative maps of axonal fibers and their density in both the native space of individual subjects and the atlas space (**Fig. 1**a).

While MAPL3 employs self-supervised pretraining with a reconstruction-based objective (**Fig. 1**c), alternative pretext tasks, including teacher-student strategies, such as DINO^45^, may offer more transferable representations. Additionally, we did not optimize the fine-tuning strategy, which has been shown to influence downstream performance^46^. In future work, we aim to incorporate such methods and extend MAPL3 to support statistical comparisons beyond atlas-defined regions, using ACE-derived^47^ local, clusterwise analysis of axonal projections across the brain. Finally, high-resolution microscopy datasets that capture time-dependent (e.g., over development, recovery from injury, treatment response, or aging) variations across axonal morphology will be used to map circuit-level perturbations with unprecedented precision.

### Ground truth label generation

Our approach for generating gold-standard ground truth (GT) labels of axonal projections consisted of two stages: classification using the LABKIT^48^ tool and post-processing refinement.

#### LABKIT classifier

To generate GT labels, we first visualized each subject’s whole-brain LSFM data as a virtual stack in FIJI/ImageJ^49,50^. Each subject’s brain regions containing axonal fibers were cropped and saved as 3D volumes. Using these data, smaller image sub-volumes (256^3^ voxels ≃ 0.46^3^*mm*^3^) were extracted using a windowing approach without overlap. To generate GT labels, we applied the LABKIT pixel classifier, which utilizes a random forest algorithm within a user-friendly interface. The random forest algorithm is a widely used supervised machine learning method that aggregates hundreds of decision trees, each trained on a slightly different set of features. Features include pixel intensity descriptors, edginess, and texture using multiple 3D filters such as Gaussian blur and Hessian eigenvalues at various scales. The final predictions of the RF are made by averaging the predictions of each tree. We used a random forest classifier with 100 trees. To train the LABKIT classifier on every 256^3^ cubes, an expert manually annotated single axons as the foreground class. The model was iteratively trained, with feedback provided by adjusting annotations for foreground and background classes and refining the predictions until satisfactory results were achieved. The final predictions of the LABKIT model were saved as a binary stack corresponding to the input image sub-volume. On average, it took approximately one hour per sub-volume, with multiple rounds of iterations per user to provide feedback to the model. That is, the initial sub-volumes would take a bit longer to be trained on the data, and as the model learns, the number of annotations and the time required per sub-volume decrease.

#### Postprocessing

To reduce the number of false positives (i.e., small or truncated objects) in the GT labels, we applied a custom postprocessing pipeline. This included a dilation operation followed by a shape filter, both of which were implemented using the ImageJ Shape Filter plugin. We first applied a dilation filter with a sphere-like kernel (radius of 1 voxel and two iterations) on the GT labels. Subsequently, a shape filter was designed and applied to the dilated image by removing objects with an area smaller than 10 pixels or when the larger side of the minimum bounding rectangle of the object was smaller than 10 pixels in each slice of the image. The final GT labels were obtained by multiplying the LABKIT-generated outputs and the post-processed image. All GT labels underwent visual quality control and correction by two independent raters (AA and SP).

From our test datasets, we randomly selected N=2112 unique patches (64^3^ *voxels* ≃0.11^3^*mm*^3^), and from our unseen dataset, we cropped N=384 unique patches of 64^3^ *voxels* 0.32^3^mm^3^, and generated manual ground truth segmentations for performance evaluation.

### Preprocessing steps

To remove the ‘stripe’ and ‘shadow’ artifacts from the LSFM datasets and increase the signal-to-noise ratio in each image sub-volume (256^3^ *voxels* ≃0.46^3^*mm*^3^), we developed a fast and computationally efficient algorithm similar to the light sheet artifact correction method previously described^51,52^. Our method leverages the fact that these artifacts occur along a predetermined axis due to the fixed orientation of the light sheet entering the sample from a consistent direction. Moreover, background voxels typically exhibit consistent intensities along the stripe axis over a specific length scale. We began by estimating the light sheet stripe artifact intensity, *lsai*, for each voxel *i*, by calculating a predefined percentile, *p* = 25, of the voxel intensities within a region centered around *i* that is highly elongated along the stripe-artifact axis. The size of this region (1 × 1 × 100 *voxels*) was selected to match the scale at which stripe artifact intensity changes, while its width and depth were constrained to be smaller than the cross-sectional area of the stripe artifacts. Subtracting this estimated artifact intensity from the image resulted in effective correction of the stripe artifacts, except at voxels associated with longer tracts aligned with the artifact axis. To address these voxels and avoid their removal, we also estimated the local background intensity, *bi*, by calculating the 25th percentile of voxel intensities in a square-shaped region centered around the voxel *i*. This region was chosen empirically to be larger than the largest axon fiber present in the brain sample. As the background estimation was time-consuming and computationally expensive when operating on a 3D image sub-volume (256^3^ voxels ≃0.46^3^*mm*^3^), we opted to downsample the image by a factor of 8, use a structuring region of 16^3^ *voxels* to estimate the background (Extended Fig. 10a), and upsample the results linearly to the same size as the image sub-volume. This method allowed us to achieve similar results as the original-resolution method for the background estimation step while being time-effective. The background estimate image was compared to the light sheet artifact estimate, and the corrected voxel intensity *ci* was obtained using the following equation:

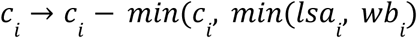

Where *w* = 2 is a factor that allows adjustment of the background estimate.

To further enhance the signal-to-noise ratio of each image sub-volume, and address the ‘blur’ or ‘stray’ artifacts that produce areas of high-intensity voxels around bright and large axon fibers, we developed a pseudo-deconvolution algorithm, as previously described^53^, with some modifications. High-intensity voxels were first identified using a computed threshold (95% percentile of the voxel intensities in each sub-volume), then smoothed with a 3D Gaussian filter with a σ = 3 *voxels*. The smoothed result was subtracted from the original image while ensuring that the values of the high-intensity voxels remained intact.

### SPECTRE network architecture

Current DL-based pipelines–mostly based on convolutional neural networks (CNNs), such as U-Net and its variants^27,54,55^–typically begin by dividing the expansive brain volumes into smaller, manageable 3D image patches, sometimes as small as 64³ voxels, within a larger sub-volume (e.g., TrailMap^2^). While CNNs are effective at capturing local morphological features, they often struggle to extract long-range dependencies and global context within these individual image patches. Vision transformers (ViTs)-based architectures^56^ offer an alternative by leveraging self-attention mechanisms and treating images as one-dimensional vectors of embedded patches to efficiently learn global contextual information. Despite this advantage, transformers may overlook local structures, and they require substantial amounts of labelled data to avoid overfitting (given their large number of parameters), a difficult and time-consuming barrier in axonal fiber segmentation in tera-voxel LSFM datasets. Even with sufficient labelled data, the prevalent patch-wise training and processing approach, employed by both CNNs and ViTs fail to exploit the inherent relationships *between adjacent image patches within a large sub-volume*, where neighbouring regions often share similar noise patterns, signal distributions, and appearance of axonal fibers. Neglecting these between-patch relationships limits the accuracy and generalizability of DL models in large-scale microscopy datasets.

We based SPECTRE on the 3D Residual U-Net^54,55^, which utilizes an encoder-decoder structure with skip connections. 3D image sub-volumes (Height × Weight × Depth; 256^3^ voxels) are used as input and reshaped to *N_Patc_*_ℎ_× 64^3^ *voxels* (*N_Patc_*_ℎ_ = 64) and processed using five blocks of encoders, consisting of three consecutive convolution layers. We have added a stochastic depth (drop path) module to each layer that randomly drops entire residual blocks during training and bypasses their transformations through skip connections (Fig.1), thus addressing the vanishing gradient and feature reuse issues in deep neural networks^57^. Parametric rectifying linear units^58^ have been deployed to provide different parameterized nonlinearity for each layer. We used batch normalization instead of instance normalization to achieve a stable distribution of activation values throughout the training and to accelerate the training^59^. A Dropout layer has been added between all convolution blocks in the architecture as a regularizer to avoid overfitting^60^.

Meanwhile, the decoder with up-convolution operators leverages the extracted feature representations from the encoder to classify all pixels at the original image resolution (256^3^ voxels). We incorporated deep supervision^61^ into the decoder, which allows a better gradient flow by computing the loss function on different decoder levels. Specifically, we used the outputs of the last three layers of the decoder to calculate the loss function. Except for the first residual units, all the convolutions and transpose convolutions had a stride of two for downsampling and upsampling. The first residual unit used a stride of one, which has been shown to improve the performance by avoiding immediate downsampling of the input image patch.

Within SPECTRE’s architecture, we introduce two sets of transformer blocks^56^ (Fig.1) that strive to capture dependencies and associations both *between* and *within* patches of LSFM datasets. The resulting features from the last layer of the encoder are passed to the globally aware block, consisting of a patch-wise transformer and a voxel-wise transformer, and a skip connection^62^, bypassing transformers and connecting the last layer of the encoder directly to the decoder. While the voxel-wise transformer is responsible for learning dependencies within each patch, our novel patch-wise transformer attends across patches and captures associations among adjacent cubic volumes. Each transformer block is conjugated with a learnable weight, allowing the model more freedom to learn and determine the relative importance of voxel- or patch-wise information, given the segmentation task. In addition, we introduce a globally aware bias term, where the resulting information from the patch-wise transformer is passed to a multi-layer perceptron to produce a bias term per patch for each channel. Each generated bias is broadcast and used to shift the final logits in the associated output volume. This bias for each image patch is learned using patch-wise information.

### Self-supervised auxiliary task

Self-supervised learning (SSL) approaches have been introduced to address the DL models’ hunger for data^63,64^. In SSL, auxiliary tasks are created to enable models to learn relevant and meaningful data representations from unlabeled datasets. These models can then be fine-tuned for segmentation tasks, reducing the reliance on exhaustive ground-truth annotations and improving model robustness on out-of-distribution datasets. Although SSL techniques have shown initial promise in microscopy^20,21,63^, current evaluations on axonal fiber mapping are limited to simple SSL frameworks and patch-level performance on small datasets. As such, existing methods lack scalability and specificity for the unique demands of 3D LSFM data, and their applicability to brain-wide connectivity mapping remains unexplored.

We introduced a reconstruction task as an auxiliary SSL task. To simplify the masking process, we employed a strategy of randomly masking 3D cubes uniformly over the foreground regions rather than using blockwise masking approaches [e.g., BEiT^65^]. The DL model predicted the raw intensity values corresponding to the masked areas. Input images were normalized by scaling their minimum and maximum intensities to 0 and 1, respectively. Random image augmentations, used in real-time at every pretraining epoch, included random contrast adjustment and intensity shift.

To implement our masking strategy, two transforms were randomly (with a probability of *P* = 0. 5) applied to each 256^3^ sub-volume at every epoch. Both transforms were based on the MONAI^66^ *RandCoarseDropoutd*. In the first one, the *dropout_holes* parameter was set to False, masking areas outside the random locations and filling them with random values between the sub-volume minimum and maximum. For this transform, 10 randomly selected 3D cubes with spatial sizes ranging from 100^3^ voxels to 250^3^ voxels were masked. The second transform was tailored to account for the large class imbalance in LSFM data, where background voxels dominate. Here, the masking locations were constrained to foreground regions, identified using a threshold of 0.5 for each normalized sub-volume. Within the foreground regions, 10 cubes were randomly masked with spatial sizes ranging from 32x32x2 voxels to 96x96x5 voxels. Masked areas were filled with random intensity values sampled uniformly from the minimum and maximum intensities of the respective sub-volumes. Parameters for each data augmentation were independently sampled at each epoch from a predefined range of values.

### Self-supervised loss function

To optimize the self-supervised auxiliary task, we employed an L1 loss function (mean absolute error) to compare the input image (*I*; 256^3^ sub-volume with masked areas) and its corresponding ground truth (*GT*; 256^3^ sub-volume without masking). To emphasize the masked regions and address the class imbalance issue, we incorporated a dynamic weighting scheme based on the epoch number (*e*) and a scaling factor (υ = 0. 5). Voxels in the masked locations were weighted using the scaler and a value of λ*masked* = (1 − υ × 0. 99^e^), with non-masked voxels weighted with a value of λ*_non_*_−*masked*_ = υ × 0. 99^e^:

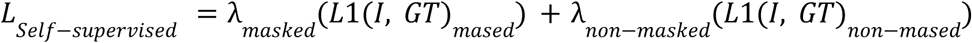

We used deep supervision^61^ in our learning strategy, where losses are computed at multiple decoder levels to improve gradient flow, defined as:

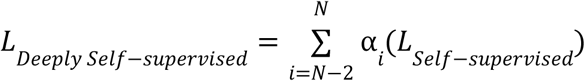

where *N* is the final decoder layer, and α*_i_* are layer-specific weights and equal to 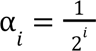 for i=0, 1, 2, pointing to the *Nt*ℎ, (*N* − 1)*t*ℎ, and (*N* − 2)*t*ℎ layer, respectively.

### Finetuning Loss function

To optimize segmentation performance, we developed a hybrid loss function that combines an edge-weighted loss, a deeply supervised DiceFocal loss^67^, and a Maximum Intensity Projection^68^ (MIP) loss. The total loss was computed as a weighted sum of these components:

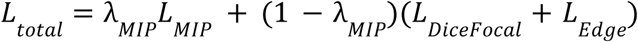

where λ*_MIP_* is a hyperparameter (set to 0.3) weighting the contribution of the MIP loss relative to the other loss functions, optimized for performance on the validation set.

#### Edge-weighted Loss

To emphasize axonal edges during training, we incorporated a *Binary Cross-Entropy* loss, weighted to prioritize edge regions:

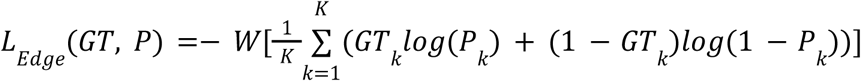

Where *K* is the number of voxels, *GT_k_* and *P_k_* denote the ground truth and the output probability map at voxel *k*, respectively, and a weight matrix (*W*), determined from the true labels. Specifically, we dilated (twice) the true labels using a max pooling operation with a kernel size of 3, and padding and stride size of 1. The resulting dilated image is then subtracted from the original ground truth map to compute the edge of the axons. Voxels in the image containing edges were weighted with a value of *Wedge* = 0. 7, with non-edge voxels weighted with a value of *Wot*ℎ*er* = 0. 3.

#### Deeply supervised DiceFocal loss

We used a DiceFocal loss function that returns the sum of Dice loss and Focal Loss. We used focal loss to address the large class imbalance encountered during training and mitigate the dominance of easily classified negatives on the cross-entropy loss and its gradients. Focal loss introduces a modulating factor (1 − *P_t_*)^γ^, (γ set to 2) to attenuate well-classified negative examples and emphasize harder, misclassified examples during training. For notational convenience, *Pt*(*t* is an index for the true class label) equals *P* if *y* = 1 and 1 − *P* otherwise, where *P* is the output probability map and *P* ∈ [0, 1]. The Focal loss is calculated as:

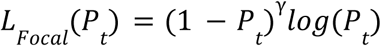

The Dice coefficient (between 0 and 1) measures the similarity between two labels, which we aimed to maximize. The Dice coefficient can be computed as:

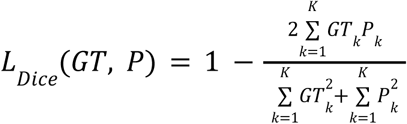

Where the sums are computed over the *K* voxels.

Deep supervision computes losses at multiple decoder levels to improve gradient flow, defined as:

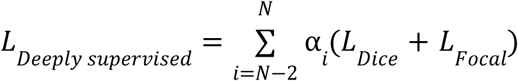

where *N* is the final decoder layer, and α*i* are layer-specific weights and equal to 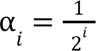 for *i* = 0, 1, 2, pointing to the *Nt*ℎ, (*N* − 1)*t*ℎ, and (*N* − 2)*t*ℎ layer, respectively.

#### Maximum Intensity Projection (MIP) Loss

To improve axonal continuity, we incorporated a loss based on Maximum Intensity Projections (MIP) along the three principal axes (*x*, *y*, and *z*). This was calculated by comparing the MIP of predicted binary segmentations with that of the ground truth using DiceFocal loss:

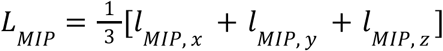

where *lMIP*, *x* , *lMIP*, y , and *lMIP*, z denote DiceFocal losses for the projections along each axis.

We selected the configuration yielding the highest Dice coefficient on the validation set by systematically varying the weights and scaling factors.

### Augmented Patch Generation

A major challenge in robust axonal segmentation in tera-voxel LSFM arises from the extreme class imbalance between axon-containing regions and background or imaging artifacts, compounded by the difficulty of generating reliable ground truth annotations. As a result, training datasets often lack enough diversity and fail to capture complex scenarios such as when axons and high-intensity artifacts co-occur within the same patch. This leads to frequent segmentation errors: some models produce widespread false positives in artifact-prone regions, while others miss true axonal signals altogether, reflecting differences in data composition and training objectives. This means that each analysis pipeline offers different perspectives and biases, ultimately hindering the accurate reconstruction of whole-brain connectivity and limiting biological interpretation.

To address this challenge, we designed an automated augmentation strategy that introduces biologically plausible axonal structures into hard negative patches (regions devoid of axonal fibers but containing challenging bright artifacts such as vasculature or neuronal somas). This augmentation enables the model to better distinguish true axons from morphologically similar false positives in regions that training patches do not cover. A total of 150 hard negative 3D sub-volumes (256^3^ voxels) were randomly extracted from regions near anatomical boundaries and structures with high-intensity non-axonal signals. For augmentation, we randomly paired each hard negative sub-volume with an axon-containing training sub-volume, and generated 460 augmented 3D sub-volumes.

Each augmented sub-volume was constructed as follows:

#### Axon Cube Extraction

From the selected axon-containing sub-volume, we identified the axonal regions using the ground truth and applied 3D binary dilation (using *binary dilation* function from *scipy.ndimage* library with a 3x3x3 cube structure and three iterations). A crop of size *H* × *W* × *D*, constrained within the dilated axonal region, was randomly selected to form the source cube.

#### Target Region Selection

Within the selected hard negative sub-volume, we identified candidate regions for replacement by thresholding the intensity above the 90th percentile to ensure realistic embedding locations inside the brain. A randomly selected location within the sub-volume was chosen, such that the source cube could be inserted without exceeding volume bounds.

#### SNR Normalization

To prevent unrealistic signal intensities, we adjusted the brightness of the axonal region such that the SNR of the resulting augmented sub-volume matched the SNR characteristics of the original axon-containing source sub-volume. Let 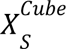 and 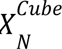 denote the voxel intensities of the axonal (signal) and non-axonal (noise) regions within the source cube, and 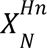 denote the background intensities at the insertion location in the hard negative sub-volume. The scaling factor α applied to the axonal signal was computed as:

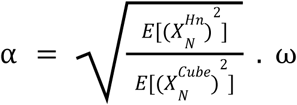

where ω ∼ *u*(0. 5, 2) is a randomly sampled jitter term to simulate natural SNR variability. The axonal voxel values were then rescaled as:

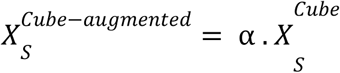

#### Patch Reconstruction

The modified axonal intensities 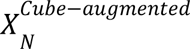 and their corresponding labels were inserted into the chosen location in the hard negative sub-volume, creating a new augmented sub-volume. This SNR-aware augmentation not only preserved the spatial and textural characteristics of real axonal fibers but also forced the model to learn discriminative features against visually similar distractors, ultimately improving segmentation fidelity in challenging contexts.

### Model Training

#### Self-Supervised Learning (SSL)

To pre-train our deep learning model, we employed 22,000 unique sub-volumes (256^3^ voxels; 90% for training and 10% for validation), excluding data augmentation, which were derived from five whole-brain LSFM datasets curated for pre-training. Given the architecture of our neural network, which extracts both within- and between-patch information from patches of 64^3^ voxels, our model encountered a total of 1,280,000 unique image patches during the SSL phase. Image patches containing a large number of background voxels (>25% of total voxels in each patch) were filtered out from the datasets. The model was trained for 90 epochs, with each epoch comprising 19,800 steps. Two model checkpoints were saved per epoch—at the midpoint and endpoint—resulting in 180 pre-trained models. To identify the most effective pre-trained model for downstream tasks, the top 30 models were shortlisted based on validation performance. These models were subsequently fine-tuned using 30% of the dataset with ground truth annotations, and the model yielding the highest validation score was selected for further supervised training (see *Supervised Learning*).

#### Supervised Learning

For fine-tuning or training from scratch, we created 256^3^ image sub-volumes from six whole-brain LSFM datasets (5 for training and validation, and one whole-brain for testing). To address the class imbalance issue in our dataset, sub-volumes containing only background, or a small number of foreground voxels (<1% of voxels inside each image sub-volume) were filtered out. Consequently, we used 233, 47, and 33 unique image sub-volumes (excluding data augmentation) for training, validation, and testing, respectively. Given the architecture of our model, this corresponds to 14,912, 3,008, and 2112 unique 64^3^ image patches for training, validation, and testing.

We trained multiple models with different hyperparameter settings, which were optimized using grid search. The hyperparameters included input image size, encoder and decoder depth, including the number of attention heads and MLP ratio, learning rate, dropout probability, kernel size, and the weights and parameters in the proposed loss function. The final model, which achieved the best Dice score performance on the validation set, consisted of a 5-layer encoder-decoder network with an initial channel size of 16 and a kernel size of 3x3x3, 8 attention heads with an MLP ratio of 4, dropout rates of 0.1 for the encoder-decoder path, and a dropout probability of 0.05 (for both patch- and voxel-wise transformers).

The deep learning model was trained for 700 epochs. To avoid overfitting, early stopping was set to 200 epochs, after which performance (Dice score) on the validation dataset did not improve. The AdamW^69^ optimizer was used with an initial learning rate of 0.0001 and a cosine scheduler with a warm-up epoch set to 20% of the total maximum epochs^70^.

#### Finetuning with augmented sub-volumes

The deep learning model was finetuned for 20 epochs using all train sub-volumes combined with the augmented sub-volumes consisting of 774 sub-volumes (256^3^ voxels) ≃ 50,000 patches (64^3^ voxels), with no significant performance difference on the validation metric compared to the model trained without augmented sub-volumes. The AdamW optimizer was used with an initial learning rate of 0.00001 and a cosine scheduler with a warm-up epoch set to 20% of the total maximum epochs.

### Postprocessing steps

Our postprocessing algorithm consisted of two stages: a thinning/skeletonization algorithm and post-hoc morphological filtering.

#### Thinning/Skeletonization algorithm

To generate a refined skeletonized representation, we processed each 2D slice of the probability map on the GPU using CUCIM (a GPU-accelerated version of scikit-image). The probability map was first binarized at nine thresholds, ranging from 0.1 to 0.9. For each binarized map, the distance transform map to the background was computed using the *medial_axis* function from the skimage.morphology Python library. The resulting distance transform maps were concatenated by a voxel-wise sum. The final skeletonized map was then derived by identifying the ridge of the concatenated distance transform using the *peak_local_max* function from the skimage.feature library. This skeletonization approach connects fragmented axonal segments that arise from threshold segmentation, producing a continuous and coherent representation of the underlying axonal structures. The procedure was run slice-by-slice in 2D with GPU acceleration and parallelized across files/slices for efficiency.

#### Morphological filtering

A custom GPU-accelerated shape-filtering script was implemented in Python to refine the thinned maps by identifying and removing small, truncated, or disconnected components. Connected components (using the *label* function from cuCIM.skimage.measure) and their morphological properties (using the *regionpropstable* function from skimage.measure) were calculated within the combined skeleton. The filter then excluded objects based on their geometric properties, including size (fewer than 64 voxels), elongation ratio (the ratio of inertia tensor eigenvalues, λ₁/λ₂ ≥ 5), and topology (objects with excessive internal holes, Euler number constraint, default ≤ 100). This targeted filtering step eliminates noise and artifacts, ensuring that only elongated and biologically plausible structures are retained in the final skeletonized representation.

### Monte Carlo dropout and uncertainty estimation

To estimate epistemic uncertainty during segmentation, dropout layers were retained in their stochastic state during inference, while all other model parameters remained fixed. Monte Carlo dropout can be interpreted as approximate Bayesian inference in a deep Gaussian process, with repeated stochastic forward passes providing samples from an approximate predictive distribution. For each input sub-volume, 50 stochastic forward passes were performed. Each pass generated a voxel-wise axonal probability map, *p_t_*(*x*), where t denotes the stochastic inference pass, and x denotes the voxel location. The final ensemble probability map was calculated as the voxel-wise mean across the 50 predictions:

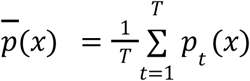

Where *T* = 50. The corresponding voxel-wise epistemic uncertainty map was calculated as the standard deviation across predictions:

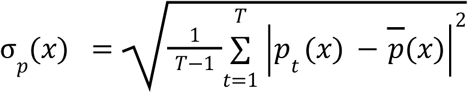

The ensemble probability map was thresholded using the same decision threshold applied during deterministic MAPL3 inference to generate the final binary segmentation. Segmentation performance was evaluated using the same precision, recall, Dice similarity coefficient, and intersection-over-union metrics used for the deterministic model. The relative change in precision was calculated as the mean absolute difference between MC-dropout and deterministic precision across unseen samples, divided by the mean deterministic MAPL3 precision.

### MAPL3 Tractography

#### Structure tensor estimation and streamline seeding

MAPL3 tractography was performed using voxelized segmentation maps registered to the ARA space at 25-µm isotropic resolution. A structure tensor field was computed after Gaussian smoothing (σ = 2.0 voxels; 50 µm) to reduce high-frequency segmentation noise while preserving directional continuity along labeled axonal bundles. Fractional anisotropy (FA) and the principal eigenvector were extracted at each signal-positive voxel to represent local fiber coherence and dominant orientation, respectively. The injection-site mask was defined using the corresponding right orbital area (RORB) subdivisions in the Allen hierarchy and dilated by 25 voxels (625 µm) to encompass perisomatic labeling and provide complete seed coverage. The segmented-signal mask was thresholded (MIN_SIG=0.1, voxels with |seg| >0.1 treated as signal) and independently dilated by 35 voxels (875 µm). Fiber orientations were propagated into the dilated signal envelope using a nearest-neighbor Euclidean distance transform, allowing streamline propagation across short discontinuities in the segmented axonal signal. Regions containing fewer than 50 signal-positive voxels were excluded from regional FA summaries to reduce instability in sparsely labeled regions. In addition to the orientation field, three tractography-independent regional image descriptors were calculated: mean FA, maximum-intensity-projection depth variance, and mean Euclidean distance from the injection site. These descriptors represented regional fiber coherence, signal dispersion along the tissue-depth axis, and anatomical distance from the injection locus, respectively.

#### Deterministic streamline reconstruction with gap tolerance

Streamlines were uniformly seeded within the dilated injection-site mask and propagated using a deterministic fiber-following algorithm. At each propagation step, the local direction was obtained from the structure tensor orientation field after Gaussian smoothing (σ = 1.0 voxel). Streamlines were advanced using a step size of 1.0 voxel (25 µm) and a maximum turning angle of 75° per step. A gap-tolerance criterion was applied to bridge discontinuities in sparsely segmented axonal labeling, allowing propagation through unlabeled voxels when supported signal could be recovered within the specified search distance (500 steps). Streamlines shorter than 1.0 mm or longer than 300.0 mm were removed. Streamlines ending outside the brain mask or within excluded atlas regions were also discarded. Of 799,906 initially generated trajectories, 726,180 passed the initial length criterion, and 623,468 remained after brain-mask and endpoint filtering (mean length, 5.09 mm; range, 1.00–32.60 mm).

#### Streamline postprocessing and density-map generation

Raw tractography output was subjected to a sequential postprocessing pipeline to remove implausible trajectories and generate quantitative spatial density maps. First, a curvature filter was applied directly to the 623,468 raw streamlines, discarding any trajectory containing a run of 0.1 mm or more with local curvature below 1.5 mm^-1^, a threshold designed to eliminate implausibly straight segments reflecting tracking artifacts rather than anatomically valid axonal paths, reflecting the expectation that genuine projections maintain measurable bending as they traverse curved white-matter architecture. The curvature-filtered streamlines were then passed through a *MIRACL spline filter* with a step length of 0.1 mm, resampling each trajectory at 0.1 mm, resampling each trajectory at uniform arc-length intervals to produce smooth, continuously parameterized streamlines stored in TRK format. Streamlines were subsequently converted to MRtrix TCK format, and a final length filter (1.0–300.0 mm) was re-applied using *tckedit* to enforce consistency across the full pipeline. Tract density index (TDI) and streamline endpoint density maps were generated in Allen CCF space at 25 µm isotropic resolution using *tckmap*, providing voxel-wise counts of streamline traversals and terminations, respectively, for visualization and quantitative regional analysis.

#### Regional traversal and streamline endpoint analysis

Each streamline was oriented from its injection-site seed toward its distal endpoint and annotated using the Allen CCF hierarchy. For every streamline, MAPL3 recorded the injection-site origin, all atlas-defined regions intersected along its trajectory, and the atlas-defined region containing its distal endpoint. Regional traversal counts were defined as the number of streamlines intersecting a given region, whereas regional endpoint counts were defined as the number of streamlines whose distal endpoints were located within that region.

To distinguish regions predominantly traversed by streamlines from regions enriched in reconstructed endpoints, we calculated the regional streamline endpoint fraction:

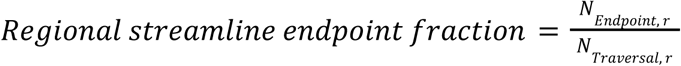

where*N_Endpoint_*, *_r_* denotes the number of distal streamline endpoints within region r, and *N_Traversal_*, *_r_* denotes the number of streamlines intersecting that region. Because every endpoint-containing streamline also intersects the corresponding endpoint region, the endpoint fraction is bounded between 0 and 1. An anatomical comparison was performed between a white-matter fiber-tract reference set and a gray-matter reference set containing established OFC projection sites. The white-matter set included the bilateral internal capsule (int), corpus callosum (cc), cerebral peduncle (cpd), cingulum bundle (cing), stria terminalis (st), columns of the fornix (fx), medial longitudinal fascicle (mlf), anterior commissure, temporal limb (act), external and internal medullary laminae (em and im), optic tract (opt), fasciculus retroflexus (fr), stria medullaris (sm), mammillothalamic tract (mtt), external capsule (ec), habenular commissure (hbc), ventral tegmental decussation (vtd), genu of the corpus callosum (ccg), and corpus callosum, posterior forceps (fp). The gray-matter set included the bilateral mediodorsal thalamic nucleus (MD), ventral tegmental area (VTA), periaqueductal gray (PAG), caudoputamen (CP), nucleus accumbens (ACB), medial amygdalar nucleus (MEA), olfactory tubercle (OT), nucleus of the diagonal band (NDB), piriform area (PIR), medial preoptic area (MPO), basolateral and central amygdalar nuclei (BLA and CEA), lateral hypothalamic area (LHA), substantia innominata (SI), midbrain reticular nucleus (MRN), posterior thalamic complex (PO), submedial thalamic nucleus (SMT), superior central nucleus raphe (CS), and ventral cochlear nucleus (VCO). The prefixes R and L denote right- and left-hemisphere structures, respectively. Each reference set initially comprised 38 bilateral regions. Because the precision of EF depends on the number of traversing streamlines, regions were included only when they met the minimum traversal threshold specified for each analysis. At the primary threshold of at least 200 traversing streamlines, 14 white-matter fiber-tract regions and 26 gray-matter regions were retained. These sets were used as anatomical reference groups rather than ground-truth classifications of individual streamlines, because white-matter tracts may contain branching or prematurely ending reconstructed streamlines, whereas gray-matter regions may contain both traversing and locally arborizing axons. The ability of the endpoint fraction to distinguish major white-matter fiber tracts from established gray-matter projection regions was evaluated using the abovementioned reference sets. Discrimination was quantified using receiver operating characteristic analysis and the area under the receiver operating characteristic curve. Differences between the reference groups were assessed using a one-sided Mann–Whitney U test. Robustness to regional sampling density was evaluated by repeating the analysis at minimum traversal-count thresholds of 50, 200, 1,000, and 2,000 streamlines. The optimal endpoint-fraction cutoff was determined using Youden’s JJJ statistic. Spearman rank correlation was used to assess the relationships of endpoint fraction with regional traversal count and distance from the injection site.

### Registration

To achieve spatial alignment between whole-brain tissue-cleared microscopy images, segmentation maps, and the Allen Mouse Brain Atlas (ARA), we employed our open-source MIRACL platform^23^. This platform is specifically optimized for the multimodal registration of cleared data, incorporating tools based on the ANTs library^71^ (http://stnava.github.io/ANTs/). The registration process involves a series of image preprocessing steps, such as denoising, correcting intensity to address brightness variations across the images, and reorienting the images to match the ARA standard orientation. Following preprocessing, an intensity-based alignment workflow consisting of two steps is performed. First, an initial rough alignment is performed using the antsAffineInitializer tool from ANTs. The second stage involves a more refined intensity-based registration using a multistage B-spline algorithm. This algorithm consists of three levels of transformation: a rigid alignment with 6 degrees of freedom (DOF) to handle basic rotations and translations, an affine alignment with 12 DOF to accommodate additional scaling and shearing, and finally, a non-rigid or deformable B-spline symmetric normalization to capture complex anatomical deformations. For the rigid and affine stages, normalized mutual information is used as the similarity metric to ensure accurate alignment between the ARA and microscopy data. For the deformable stage, cross-correlation is employed to enhance precision in matching finer details. The final result is a set of deformations that enable bidirectional warping between the native space of tissue-cleared microscopy images and the ARA reference space.

### Voxelization and Warping

#### Voxelization

To avoid loss of information during downsampling and warping, we voxelized the segmentation results into the 10 μm Allen Reference Atlas (ARA) space. Voxelization involved convolving the high-resolution segmentations with a spherical kernel of ∼5 μm radius—chosen to match the downsampling ratio and spatial scale of the ARA^23,47^. Within each sphere, the local average of segmented labels (e.g., cells or nuclei) was computed, producing a smoothed, voxel-based representation. This 3D map was generated using the *miracl seg voxelize* command, which is based on the *skimage* library in Python, with parallelization via *joblib* and *multiprocessing* to enhance scalability.

#### Warping

Each voxelized segmentation map was then non-linearly warped to the 10 μm ARA space using the *warp_clar* command from the MIRACL registration toolkit, applying subject-specific transformation matrices obtained from earlier registration steps.

### Model Evaluation

#### Evaluation Metrics

We evaluated the segmentation model using volume- and shape-based metrics, including the Dice similarity coefficient (DSC), recall, precision, F1-score, and the 95th percentile Hausdorff distance. The DSC quantifies the overlap between GT and P, normalized by the sum of the elements in each dataset, serving as a robust metric for measuring spatial agreement:

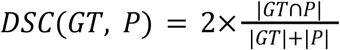

We employed the 95th percentile Hausdorff distance to evaluate spatial accuracy, a robust measure of the closeness between two point sets. Unlike the maximum Hausdorff distance, which can be sensitive to outliers, the 95th percentile provides a more representative estimate of segmentation performance. For two sets X and Y, the Hausdorff distance was defined as:

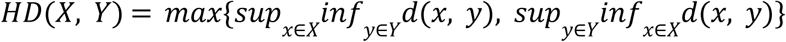

where *d*(*x*, *y*) is the Euclidean distance between points *x* and *y*.

#### Comparison against a vision transformer architecture

We benchmarked our pipeline against a vision transformer (ViT)-based segmentation model, leveraging the UNETR architecture^16,47^. UNETR replaces the encoder path of a traditional U-Net with a stack of transformer blocks to capture long-range dependencies and contextual information from embedded image patches. For this comparison, we employed an optimized UNETR variant featuring residual blocks and dropout layers to enhance robustness and generalizability. The UNETR encoder consisted of 12 transformer blocks operating on 3D input patches (*xεR*^64×64×64×1^) represented as 1D sequences of patch embeddings (16^3^ = 4096). The patch embeddings were projected into a lower-dimensional latent space (*K* = 768) and supplemented with learnable positional embeddings to preserve the spatial information. The transformer blocks integrated multi-head self-attention mechanisms and multi-layer perceptron layers to model relationships between patches. Each transformer block operates on the embedded patch sequence with the dimension of 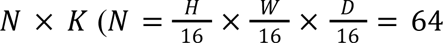 is the sequence length). Representations learned at multiple depths were reshaped and passed to the decoder via skip connections for segmentation output. Predictions were generated as 3D volumes containing voxel-wise probabilities of neuronal cell bodies (0 < *p* < 1), which were binarized using a threshold (default = 0.5). The model was trained for 700 epochs with an early stopping flag set to 200 epochs, where performance (Dice similarity coefficient) on the validation dataset did not improve, to avoid overfitting. The Adam optimizer was used with an initial learning rate of 0.0001. Lastly, the best-trained model based on validation error was used to generate whole-brain segmentation maps.

#### Comparison against D-LMBmap and TrailMap

D-LMBmap^1^ segmentation maps were generated using the *Jointly.model* available in their repository (https://github.com/lmbneuron/D-LMBmap). Axonal fiber segmentation was performed on each 2D slice of 3D image patches from test and unseen datasets using the *Toolbox.ipynb* and *Axonal-semantic-segmenter/eval.py* scripts. Resulting probability maps were binarized (with a 0.5 threshold). This threshold was applied to both the test and unseen datasets.

Similarly, axonal segmentation was performed using the 3D TrailMap^2^ pipeline with the pre-trained model provided in their repository (https://github.com/AlbertPun/TRAILMAP). Test and unseen cubes were converted to 2D slices and processed using the *segment_brain_batch.py* script. As with D-LMBmap, probability maps were binarized using thresholds between 0.1 and 0.6, with 0.1 yielding optimal validation performance.

#### Fine-tuning TrailMap for axonal fiber segmentation

To adapt the pre-trained TrailMap model to our dataset, we fine-tuned it using the original training pipeline provided in its repository. Since TrailMap requires a four-class labelling scheme (background, axon, artifact, and edge), we converted our binary MAPL3 ground-truth masks into this format. Specifically, axon voxels were assigned class 2, background voxels class 1, edge voxels (8-connected XY-plane neighbours of axon voxels that are not themselves axon) class 4, and artifact voxels (pure-background voxels exceeding mean +2σ of the background intensity distribution per volume) class 3. From each of the 694 training 256³ patches, four non-overlapping 64³ crops were randomly extracted and written directly into TrailMap’s training-set directory, yielding 2,776 training crops in total. Validation crops were similarly extracted from 21 validation patches at five crops per patch (105 crops total). Fine-tuning was performed starting from the publicly available TrailMap pre-trained weights, using the unmodified train.py script with a batch size of 6, 120 gradient steps per epoch, and 30 validation steps per epoch for 1,000 epochs. The Adam optimizer and binary cross-entropy loss were used as in the original implementation. Training loss decreased smoothly throughout fine-tuning without signs of instability or overfitting, and the best-performing checkpoint on the validation set was selected for inference. For inference on test and unseen 3D patches, each volume was decomposed into 2D axial slices, processed with the segment_brain_batch.py script, and the resulting 2D probability slices were reassembled into a 3D volume and binarized at a threshold of 0.5.

#### Fine-tuning D-LMBmap for axonal fiber segmentation

To adapt D-LMBmap to our dataset, we fine-tuned it using the nnU-Net-based training framework provided in the authors’ repository, following the documentation they supply for custom data. D-LMBmap is built on top of nnU-Net and uses a custom attention U-Net trainer MyTrainerAxial) operating on 2D axial slices within the nnU-Net 3d_fullres configuration. D-LMBmap requires four data components for fine-tuning: axon volumes, artifacts, rough labels, and fine labels. Rough labels were the binary ground-truth masks, fine labels were the skeletonized versions of the rough labels, and artifact voxels were extracted from the rough labels using the same formula described for TrailMap above. One crop was randomly extracted from each of the training axon volumes, artifacts, and rough labels in the training set; five crops (150^3^ voxels) were obtained from each of the test samples. The labels were then skeletonized using the Toolbox.ipynb notebook from the D-LMBmap repository, producing the fine labels. The fine labels and rough labels served as ground truth for training and validation, respectively. Next, the same data augmentation techniques applied in D-LMBmap were applied to the axon volumes and artifacts. These techniques were executed through the create_data.py script provided by the D-LMBmap authors. In the end, there were 3429 training samples. The *plan_and_preprocess* command in nnU-Net was run to assemble the dataset for training, including automatic configuration of model parameters and standardization of image resolution and intensity. The model weights were initialized from the publicly available pre-trained D-LMBmap model (jointly_state_dict.pth). Five-fold cross-validation was performed over 100 epochs per fold, using Adam optimization with an initial learning rate of 3×10^-4^ and a ReduceLROnPlateau scheduler that halved the learning rate upon validation loss plateau, as implemented in the authors’ *nnUNetTrainerV2.py*. Training loss decreased smoothly across all five folds without instability, indicating consistent convergence. The best-performing fold was selected based on the highest mean Dice score on its held-out validation set, and the corresponding model weights were used for all subsequent inference. For inference on test and unseen 3D patches, each volume was decomposed into 2D axial slices and processed through the trained model; the resulting 2D probability slices were then reassembled into a 3D volume, normalized by the maximum probability value, and binarized at a threshold of 0.5.

### Implementation

All DL models were implemented in Python using PyTorch, the Medical Open Network for Artificial Intelligence (MONAI^66^), and Dino^24^ frameworks. Training and analysis were performed on the Sunnybrook Research Institute (SRI) high-performance computing servers, utilizing NVIDIA A100-SXM4 (80GB memory) and NVIDIA RTX A6000 (49 GB), and the Narval high-performance computing cluster provided by the Digital Research Alliance of Canada (www.alliancecan.ca), utilizing A100 GPUs (40 GB memory).

Data were registered to the Allen Reference Atlas (ARA) using MIRACL^23,47^, our in-house, open-source software. MIRACL is fully containerized and available as Docker and Singularity/Apptainer images (https://miracl.readthedocs.io/). Visualization of results was carried out using open-source tools and Python libraries, including Matplotlib, Seaborn, Brainrender, Fiji/ImageJ^49^, itk-SNAP^72^, and Freeview (http://surfer.nmr.mgh.harvard.edu/). Figure creation for the manuscript was facilitated using BioRender (https://www.biorender.com/).

## Notes

### Competing Interest Statement

The authors have declared no competing interest.

### Summary of Updates

This substantially revised version expands the manuscript. Major updates include controlled comparisons with fine-tuned state-of-the-art methods (TrailMap and D-LMBmap), extensive out-of-distribution evaluations, and expanded ablation studies. We introduce a brain-wide tractography module for reconstructing neuronal projections and quantifying regional streamline traversal and termination patterns. Additional improvements include Monte Carlo dropout-based uncertainty estimation, ensemble inference, and a dedicated graphical user interface with expanded documentation. The Introduction, Results, Methods, and Discussion have been substantially revised to strengthen methodological rigor, biological interpretation, and accessibility.

