## Extended Data for "Quantitative profiling of whole-brain projectomes at single-axon resolution using deep learning and high-resolution light sheet microscopy"

### MAPL3 Manuscript Extended Data

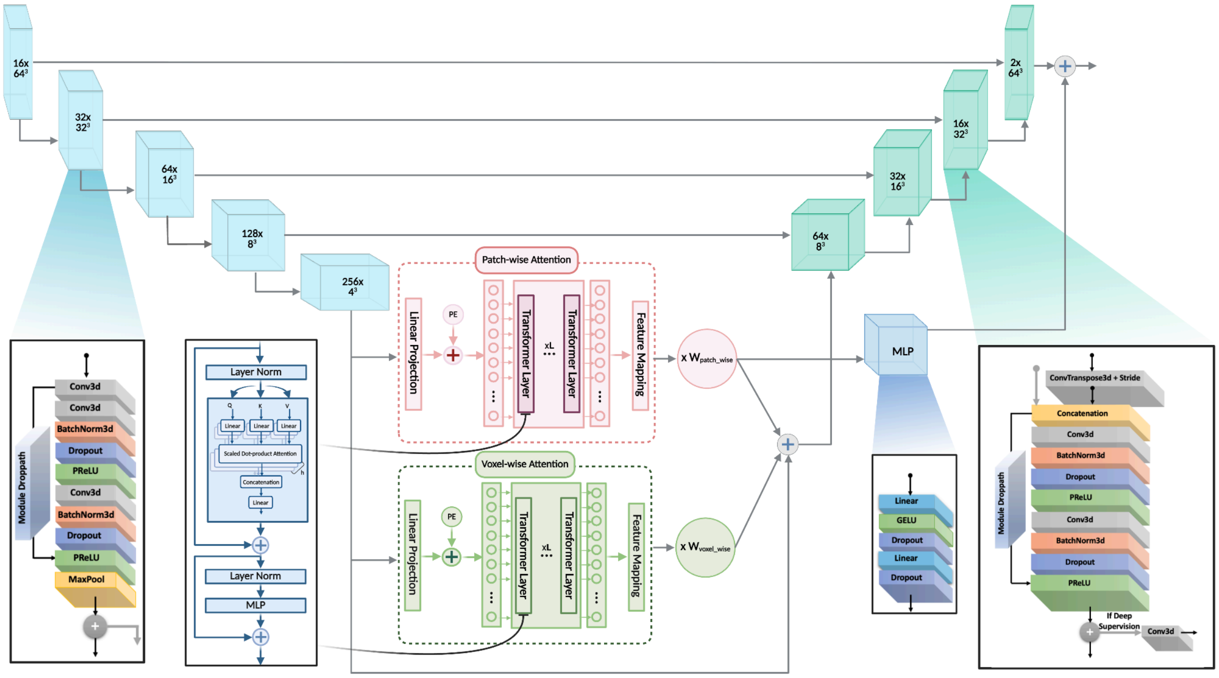

**Extended Data Fig. 1. SPECTRE Network Architecture.** SPECTRE combines a CNN with a transformer at the deepest layer, which incorporates both patch-wise (pink) and voxel-wise (green) transformer blocks with attention. Light blue boxes represent the residual encoder modules, and turquoise boxes represent the decoder modules of the CNN, with feature sizes and downsampling factors shown on the corresponding boxes. The transformer layer is shown in blue, consisting of a stack of transformer blocks, each with multi-head self-attention (8 heads, scaled dot-product attention with dropout of 0.05), residual connections with stochastic depth, and an MLP with dropout. The output of patch-wise transformer blocks is broadcast and added as an adaptive bias to the output logits. Patch and voxel-wise attention are combined using corresponding weights and fed into the CNN decoder to upsample the features back to the original space.

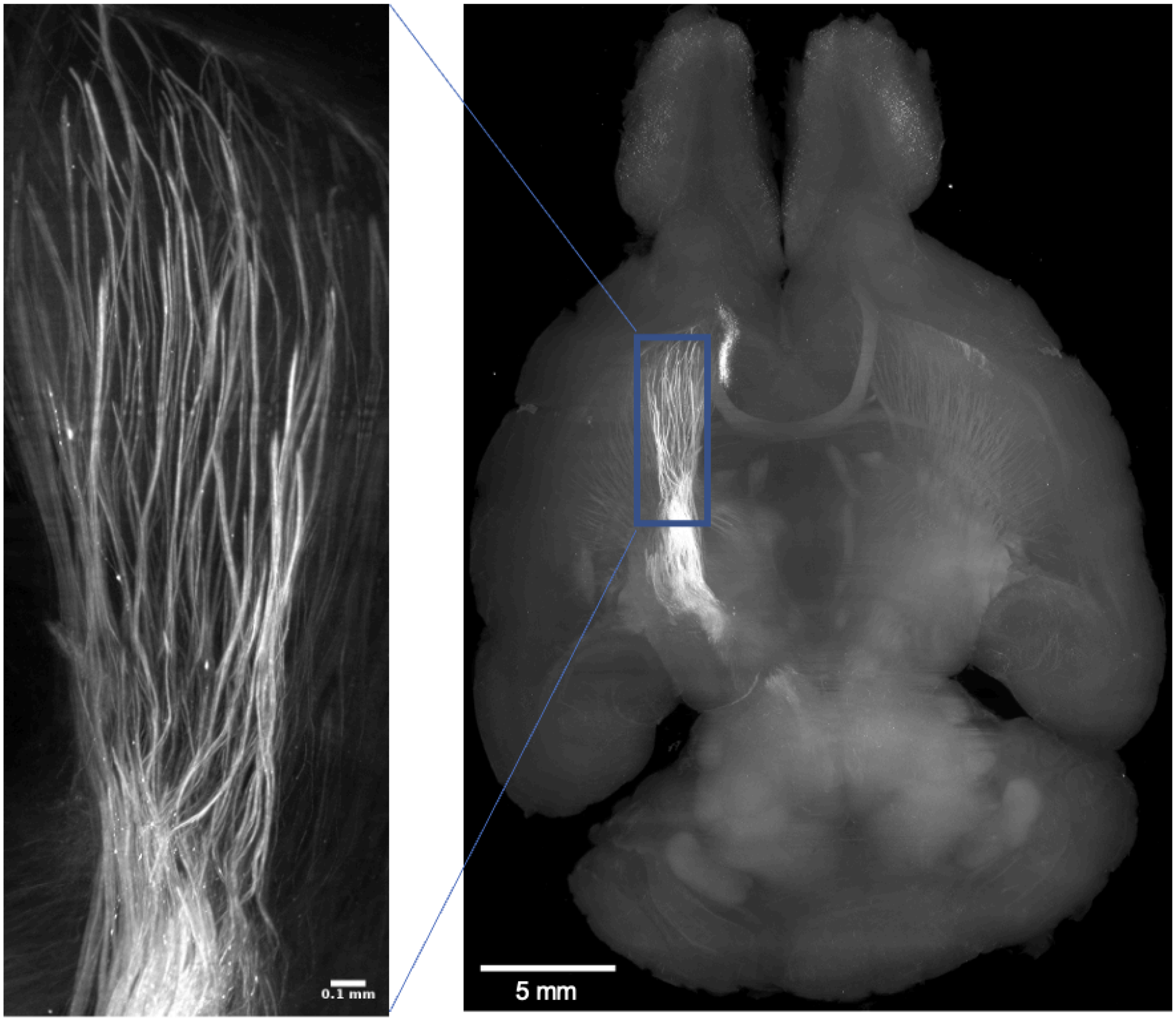


**Extended Data Fig. 2.** Axial view of whole-brain axonal projections from the orbitofrontal cortex (OFC), labeled with AAV8-CaMKIIa-eYFP-NRN, imaged on a LifeCanvas SmartSPIM (eYFP channel, 1.8 µm XY / 3 µm Z step, 3.6× objective).

a

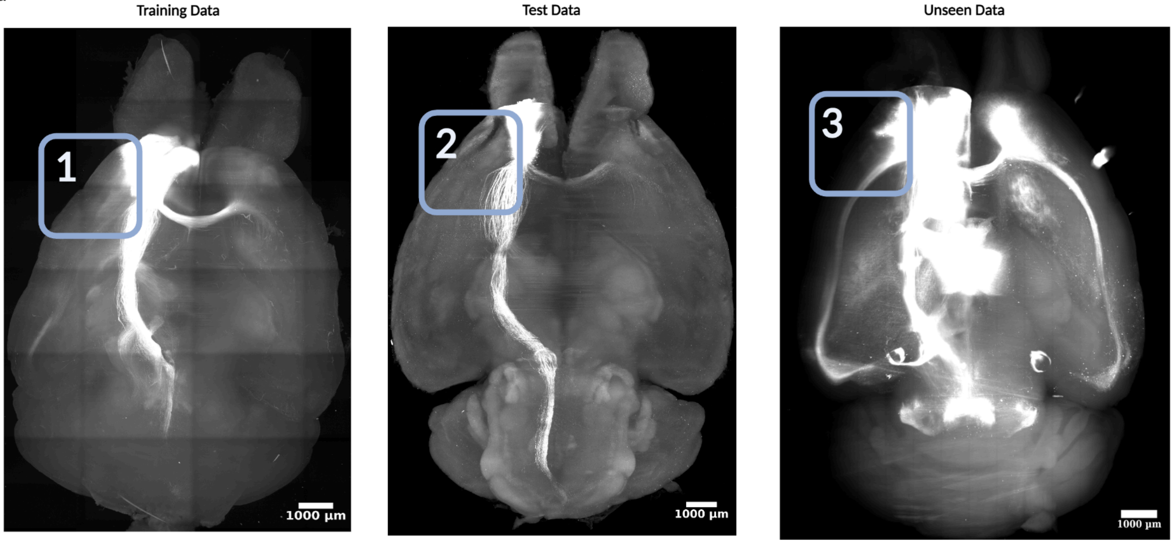


b

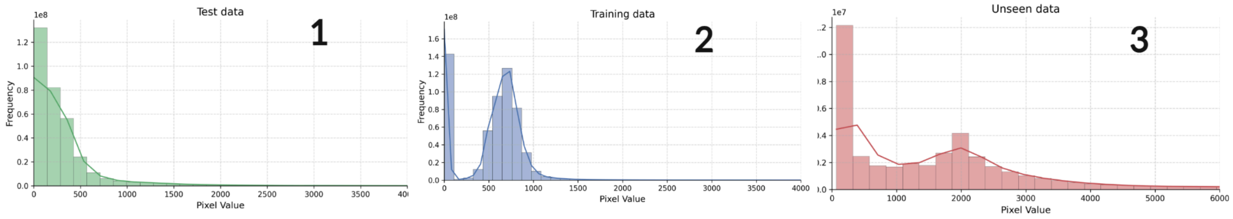


c

| | Microscope | Tissue Clearing | Voxel Size (µm) | Fluorophore | SNR (db) | CNR (db) |
|---|---|---|---|---|---|---|
| Training Data | SmartSPIM | SHIELD | 1.8x1.8x2 | eYFP | 23 ± 3 | 6 ± 1 |
| Test Data | SmartSPIM | SHIELD | 1.8x1.8x2 | eYFP | 22 ± 4 | 8 ± 1 |
| Unseen Data | LaVision | SHIELD | 5x5x5 | eYFP | 10 ± 2 | 3 ± 3 |

**Extended Data Fig. 3. Light sheet fluorescence microscopy (LSFM) datasets used to train and evaluate MAPL3.**

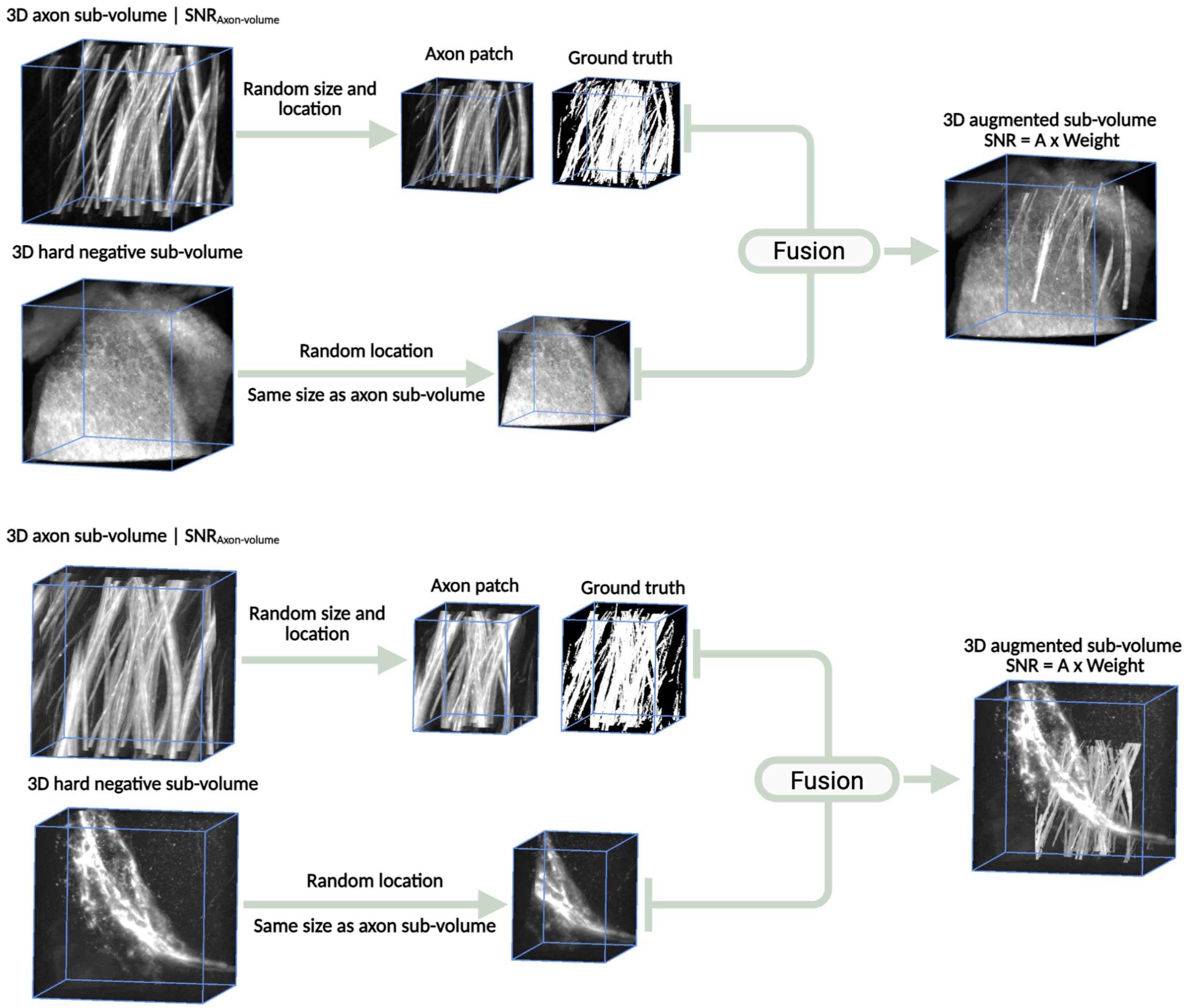


**Extended Data Fig. 4. MAPL3's patch augmentation strategy.** An automated sub-volume augmentation approach was developed to improve axon detection in challenging regions. Hard negative sub-volumes (devoid of axons but containing bright non-axonal structures such as vasculature and somas) were paired with axon-containing sub-volumes. Axonal cubes extracted from ground truth regions were embedded into hard negatives after SNR-aware intensity normalization, producing realistic augmented patches. This strategy generated 460 augmented sub-volumes and improved the model's ability to distinguish axons from morphologically similar false positives.

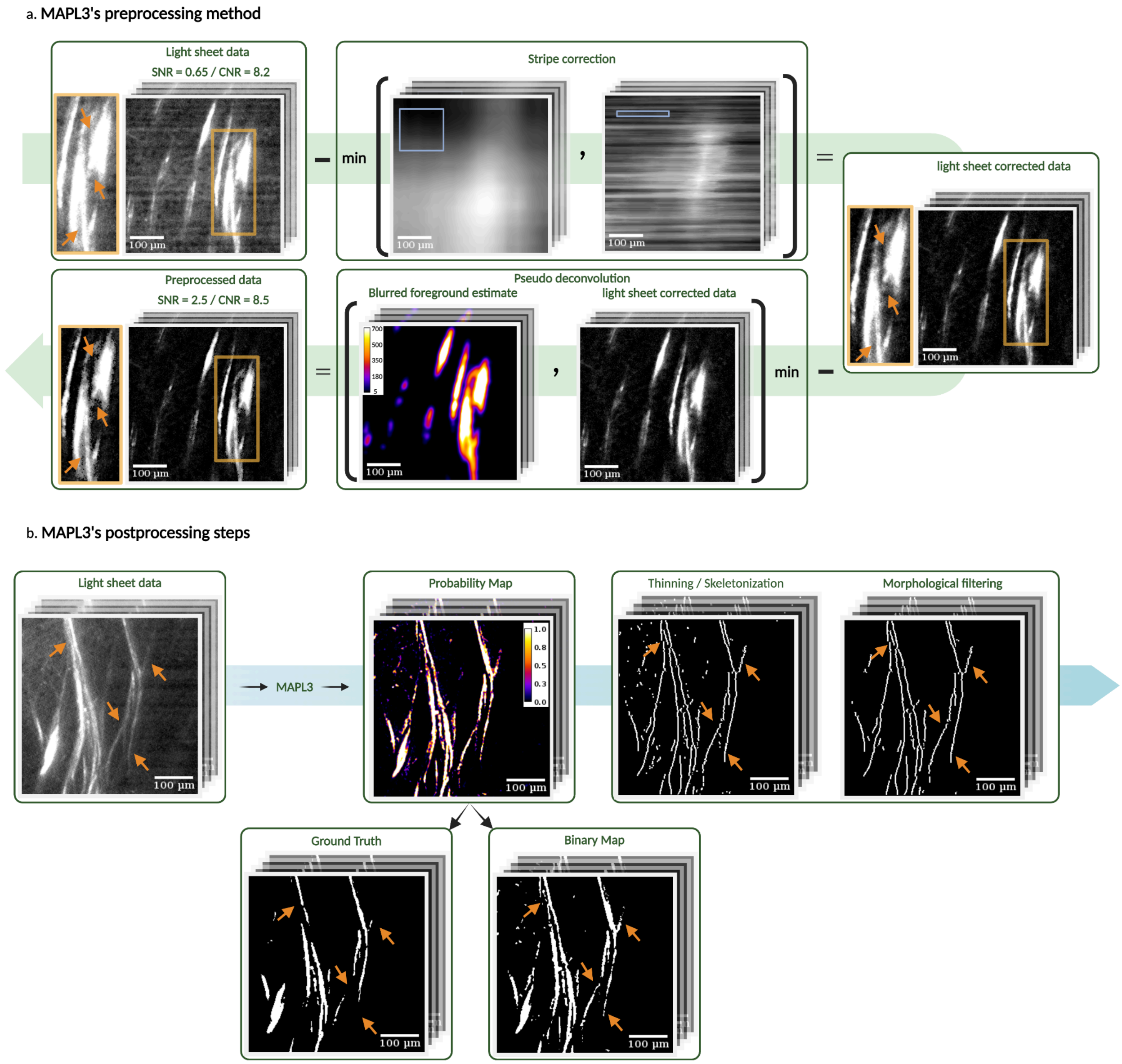


**Extended Data Fig. 5. MAPL3's pre- (a) and post-processing (b) workflows.** Pre-processing removes stripe, shadow, and blur artifacts from LSFM sub-volumes using a fast artifact-correction and pseudo-deconvolution algorithm to improve signal-to-noise ratio. Post-processing refines the segmentation outputs through multi-threshold skeletonization and morphological filtering to produce continuous, biologically plausible axonal structures.

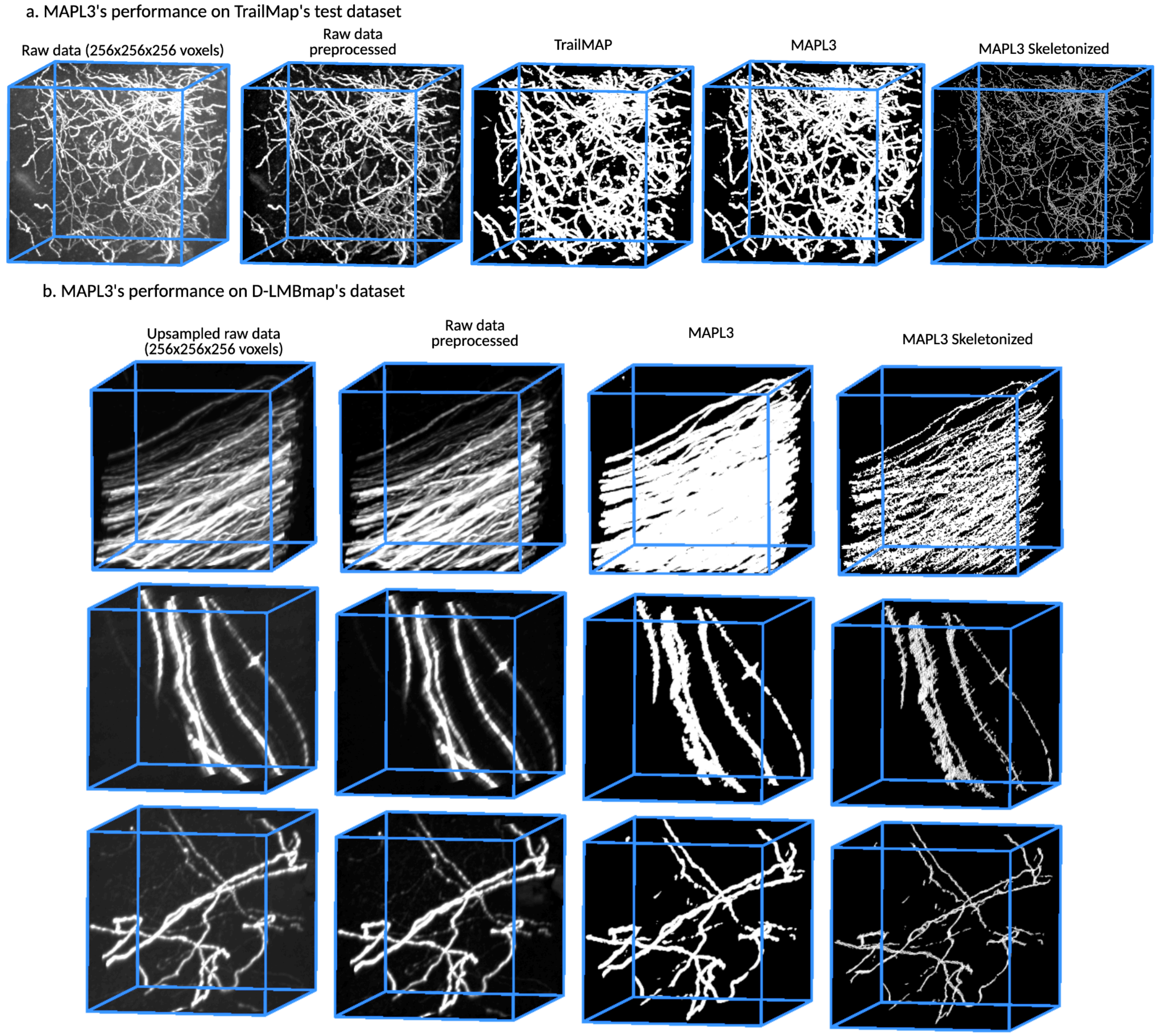


**Extended Data Fig. 6. Generalization of MAPL3 to the TrailMap and D-LMBmap benchmark datasets. a.** SPECTRE was applied to the TrailMap test dataset without fine-tuning or morphological filtering. This dataset differs substantially from the MAPL3 training data in experimental design, viral tracer, imaging protocol, and image characteristics. A binarization threshold was selected based on the predicted probability maps to generate the final segmentation, followed by skeletonization. **b.** SPECTRE was additionally applied without fine-tuning to independently acquired images from the D-LMBmap DCN-Stanford dataset. Representative sub-volumes are shown by row from the brainstem (BS), cerebellum (CB), caudoputamen (CP), cerebral cortex (CTX), and an anatomically undetermined region, respectively. MAPL3 detected axonal structures across independent datasets, anatomical regions, labeling strategies, and imaging conditions, supporting its generalizability to out-of-distribution LSFM data.

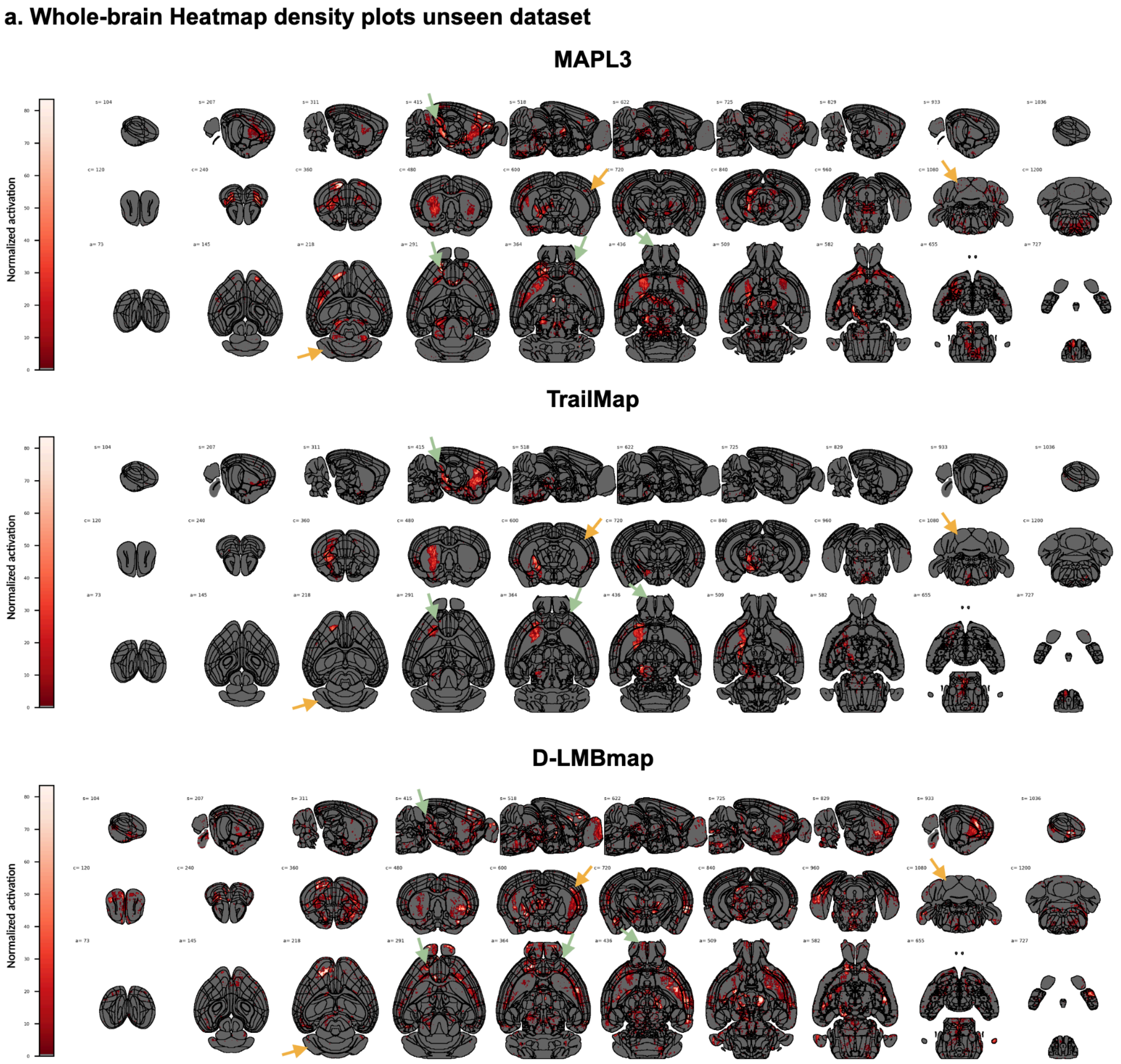


**Extended Data Fig. 7.** Whole-brain axonal projections from the right orbitofrontal cortex of an example low-resolution unseen dataset (5µm isotropic, scanned on a LaVision LSFM), reconstructed by MAPL3, TrailMap, and D-LMBmap. Voxelized segmentation maps were warped to the Allen atlas (10 µm), and region boundaries were overlaid for comparison. Rows 1–3 show sagittal, axial, and coronal views at randomly chosen cut planes. Green arrows highlight MAPL3's superior performance in detecting axonal projections and avoiding false positives, while yellow arrows show false positives in MAPL3's predictions and their comparison against TrailMap and D-LMBmap.

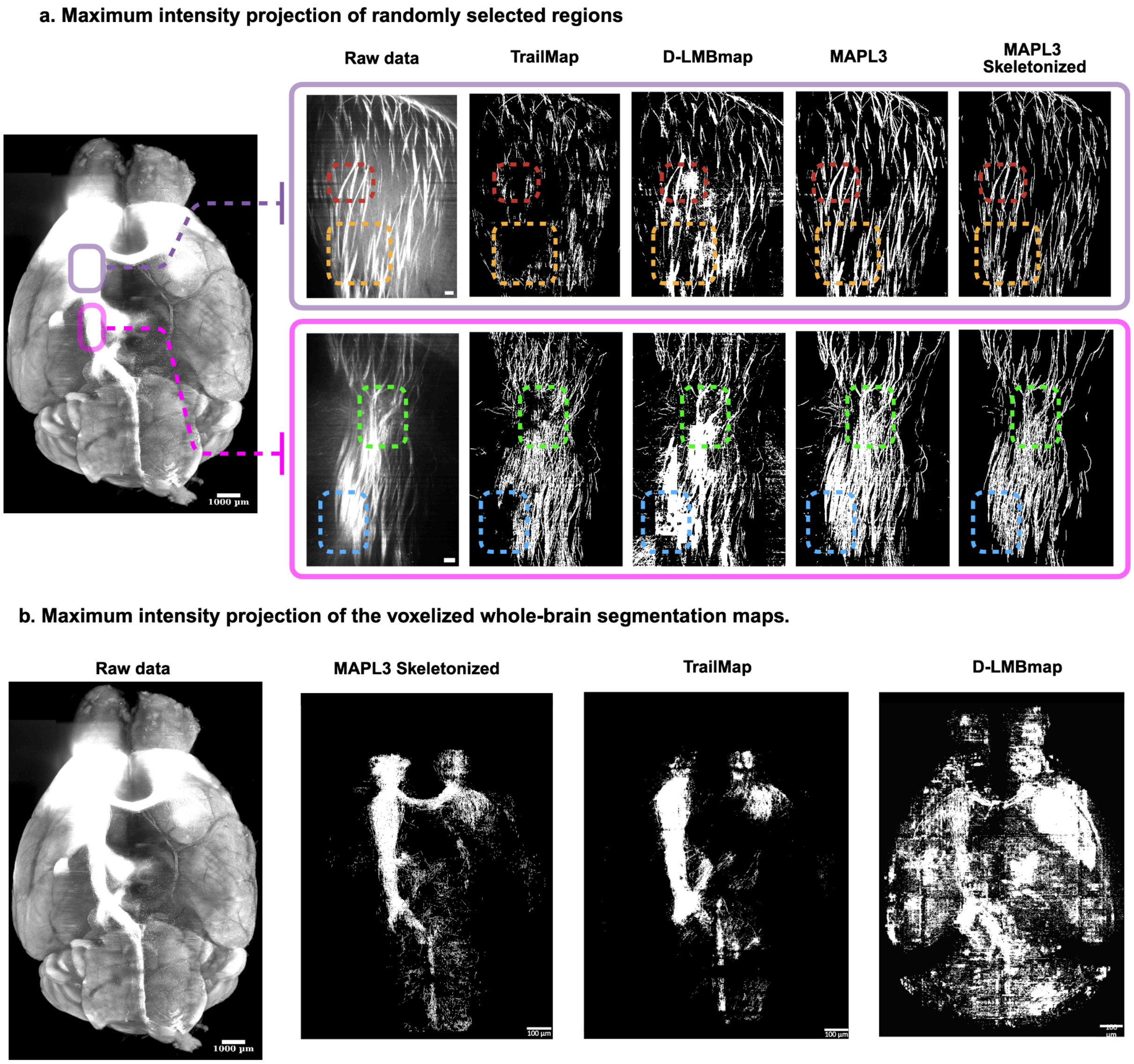


**Extended Data Fig. 8. a.** Qualitative comparison of MAPL3's segmentation module vs. TrailMap and D-LMBmap on the test dataset. Boxes highlight MAPL3 superiority in capturing axonal details compared to TrailMap and D-LMBmap. b. An axial view of the maximum intensity projection of the voxelized whole-brain segmentation maps.

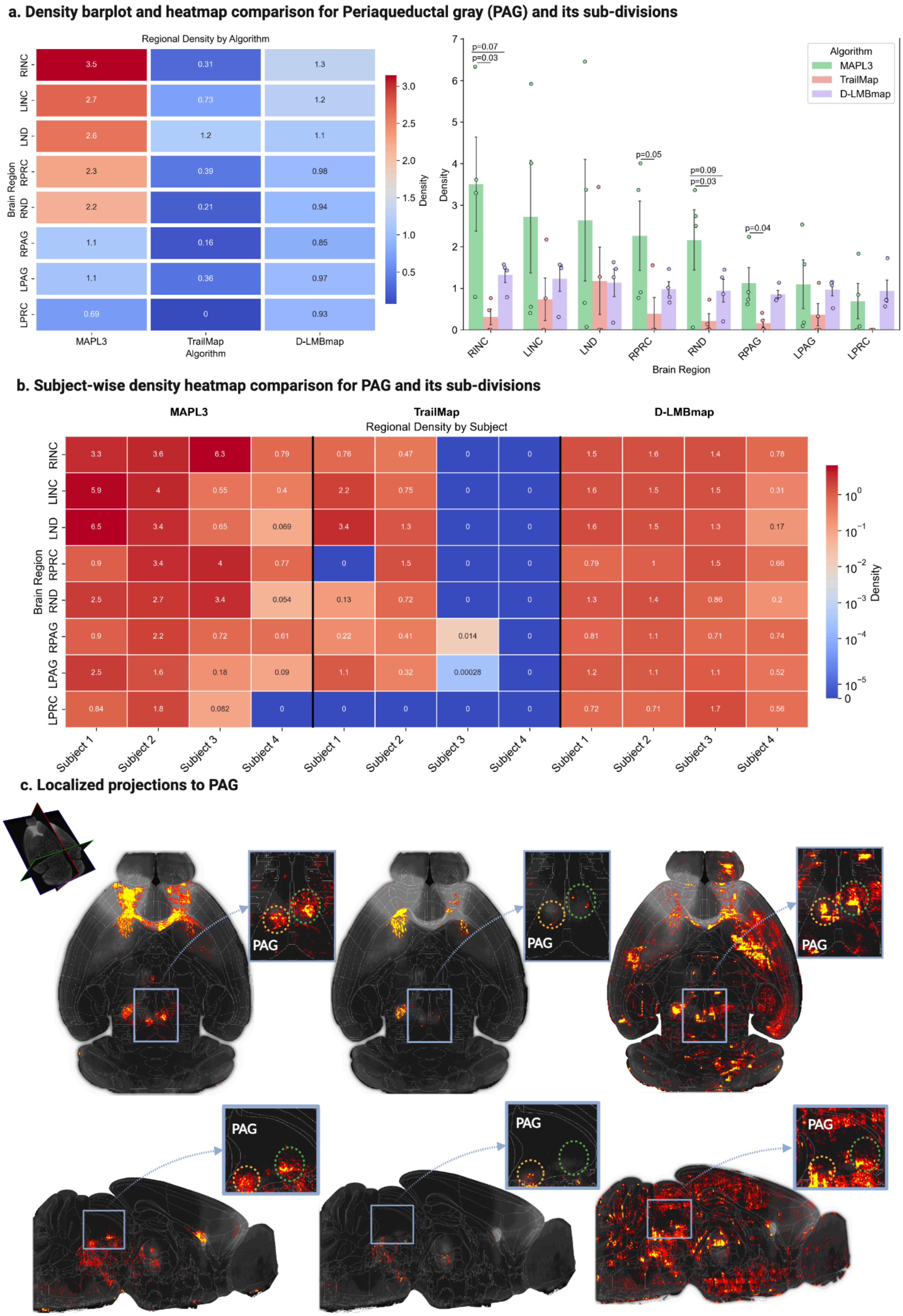
a. Density barplot and heatmap comparison for Periaqueductal gray (PAG) and its sub-divisions
Regional Density by Algorithm
RINC
LINC
LND
RPRC
RND
RPAG
LPAG
LPRC
Brain Region
3.5
0.31
1.3
2.7
0.73
1.2
2.6
1.2
1.1
2.3
0.39
0.98
2.2
0.21
0.94
1.1
0.16
0.85
1.1
0.36
0.97
0.69
0
0.93
MAPL3
TrailMap
D-LMBmap
Algorithm
Density
3.0
2.5
2.0
1.5
1.0
0.5
p=0.07
p=0.03
p=0.05
p=0.09
p=0.03
p=0.04
Algorithm
MAPL3
TrailMap
D-LMBmap
Density
Brain Region
b. Subject-wise density heatmap comparison for PAG and its sub-divisions
MAPL3
TrailMap
D-LMBmap
Regional Density by Subject
Brain Region
3.3
3.6
6.3
0.79
0.76
0.47
0
0
1.5
1.6
1.4
0.78
5.9
4
0.55
0.4
2.2
0.75
0
0
1.6
1.5
1.5
0.31
6.5
3.4
0.65
0.069
3.4
1.3
0
0
1.6
1.5
1.3
0.17
0.9
3.4
4
0.77
0
1.5
0
0
0.79
1
1.5
0.66
2.5
2.7
3.4
0.054
0.13
0.72
0
0
1.3
1.4
0.86
0.2
0.9
2.2
0.72
0.61
0.22
0.41
0.014
0
0.81
1.1
0.71
0.74
2.5
1.6
0.18
0.09
1.1
0.32
0.00028
0
1.2
1.1
1.1
0.52
0.84
1.8
0.082
0
0
0
0
0
0.72
0.71
1.7
0.56
Subject 1
Subject 2
Subject 3
Subject 4
Density
c. Localized projections to PAG
PAG

**Extended Data Fig. 9. Localized Axonal density analysis in the periaqueductal gray (PAG).** (a) Axonal fiber density within the PAG was normalized to whole-brain density for each subject, then averaged and visualized as a heatmap (left) and bar plot (right). MAPL3 reveals a smooth gradient across PAG subdivisions, reflecting localized subregional activations, whereas TrailMap and D-LMBmap exhibit substantial false negatives and false positives, respectively. **(b)** Axonal density across individual subjects highlights inter-subject variability. D-LMBmap fluctuates around unity, demonstrating over-segmentation, while TrailMap shows many false negatives, particularly in challenging datasets (subjects #3 and #4). **(c)** MAPL3 resolves fine subregional collaterals, notably within the ventrolateral PAG, while TrailMap misses these collaterals and D-LMBmap oversegments them. Axial and sagittal views of warped raw data, voxelized and warped segmentation maps from a randomly selected subject, are shown with zoomed views highlighting the PAG.

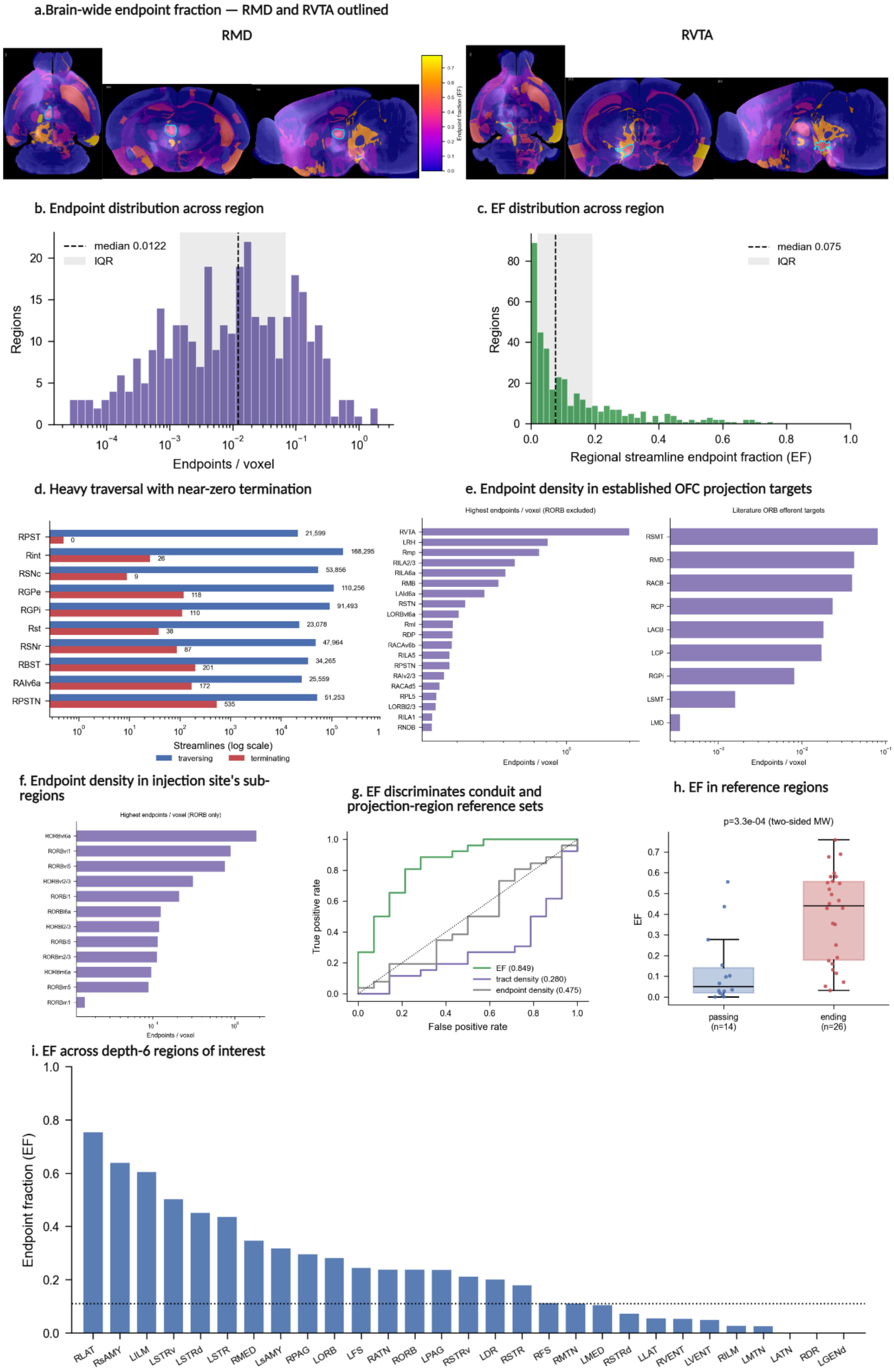

a.Brain-wide endpoint fraction — RMD and RVTA outlined
RMD
RVTA
Endpoint fraction (EF)
b. Endpoint distribution across region
median 0.0122
IQR
Regions
Endpoints / voxel
c. EF distribution across region
median 0.075
IQR
Regions
Regional streamline endpoint fraction (EF)
d. Heavy traversal with near-zero termination
RPST
Rint
RSNc
RGPe
RGPi
Rst
RSNr
RBST
RAIv6a
RPSTN
21,599
0
168,295
26
53,856
9
110,256
118
91,493
110
23,078
38
47,964
87
34,265
201
25,559
172
51,253
535
Streamlines (log scale)
traversing
terminating
e. Endpoint density in established OFC projection targets
Highest endpoints / voxel (RORB excluded)
RVTA
LRH
Rmp
RILA2/3
RILA6a
RMB
LAId6a
RSTN
LORBvl6a
RmI
RDP
RACAv6b
RILA5
RPSTN
RAIv2/3
RACAd5
RPL5
LORBl2/3
RILA1
RNDB
Endpoints / voxel
Literature ORB efferent targets
RSMT
RMD
RACB
RCP
LACB
LCP
RGPi
LSMT
LMD
Endpoints / voxel
f. Endpoint density in injection site's sub-regions
Highest endpoints / voxel (RORB only)
RORBvl6a
RORBvl1
RORBvl5
RORBvl2/3
RORBl1
RORBl6a
RORBl2/3
RORBl5
RORBm2/3
RORBm6a
RORBm5
RORBm1
Endpoints / voxel
g. EF discriminates conduit and projection-region reference sets
True positive rate
False positive rate
EF (0.849)
tract density (0.280)
endpoint density (0.475)
h. EF in reference regions
p=3.3e-04 (two-sided MW)
EF
passing (n=14)
ending (n=26)
i. EF across depth-6 regions of interest
Endpoint fraction (EF)
RLAT
RsAMY
LILM
LSTRv
LSTRd
LSTR
RMED
LsAMY
RPAG
LORB
LFS
RATN
RORB
LPAG
RSTRv
LDR
RSTR
RFS
RMTN
LMED
RSTRd
LLAT
RVENT
LVENT
RILM
LMTN
LATN
RDR
LGENd

**Extended Data Fig. 10. Brain-wide regional streamline endpoint analysis. a)** Brain-wide maps of the regional streamline endpoint fraction (EF) displayed in horizontal, coronal, and sagittal views. We also overlaid downsampled and warped Allen signal data on each panel. EF was calculated as the number of streamlines with distal reconstructed endpoints within a region divided by the number of streamlines traversing that region. The right mediodorsal nucleus of the thalamus (RMD) and right ventral tegmental area (RVTA) are outlined in cyan as representative endpoint-enriched regions. **b)** Distribution of streamline endpoint density across atlas-defined regions. Endpoint density was calculated as the number of reconstructed endpoints per voxel. **c)** Distribution of the regional streamline endpoint fraction (EF) across atlas-defined regions meeting the minimum traversal threshold. EF was calculated as the number of streamlines with distal reconstructed endpoints within a region divided by the number of streamlines traversing that region. The dashed line indicates the median EF of 0.075, and the shaded area indicates the interquartile range. **d)** Number of traversing streamlines and distal reconstructed endpoints in representative structures characterized by high traversal counts but low EF. Streamline counts are displayed on a logarithmic scale. **e)** Regional endpoint densities outside the RORB injection site. Left, regions with the highest endpoint density per voxel after excluding RORB subdivisions. Right, endpoint densities within literature-supported OFC projection regions. **f**) Endpoint density within atlas-defined subdivisions of the RORB injection site. Regions are ranked according to the number of reconstructed endpoints per voxel. **g)** Receiver operating characteristic curves comparing EF, tract density per voxel, and distal streamline endpoint density for differentiating the curated white-matter conduit and gray-matter OFC projection-region reference sets at a minimum threshold of 200 traversing streamlines. EF achieved an area under the receiver operating characteristic curve of 0.849, compared with 0.280 for tract density and 0.475 for endpoint density. **h**) EF values in the curated white-matter fiber-tracts and gray-matter OFC projection-region reference sets at the minimum threshold of 200 traversing streamlines. Points represent individual atlas-defined regions; boxes indicate the median and interquartile range, and whiskers extend to the most extreme observations within 1.5 times the interquartile range. Statistical significance was assessed using a two-sided Mann-Whitney U test ($P = 3.3 \times 10^{-4}$; 14 conduit regions and 26 projection regions). **i)** EF across selected ARA depth-6 regions of interest. The dotted line indicates the EF cutoff of 0.11 that provided the greatest separation between the two reference sets in this dataset. ARA, Allen Reference Atlas; AUC, area under the receiver operating characteristic curve; EF, regional streamline endpoint fraction; OFC, orbitofrontal cortex; ROC, receiver operating characteristic.
