## Supplementary Material for "Quantitative profiling of whole-brain projectomes at single-axon resolution using deep learning and high-resolution light sheet microscopy"

### MAPL3 Manuscript Supplementary Materials

**Supplementary Table 1.** SPECTRE performance comparison against UNETR

| Algorithm | DSC | HD95 | Recall | Precision |
|---|---|---|---|---|
| In-distribution test dataset | | | | |
| UNETR | 0.69 ± 0.03 | 4.27 ± 1.56 | 0.87 ± 0.10 | 0.59 ± 0.09 |
| MAPL3 | **0.74 ± 0.04** | **4.27 ± 1.73** | **0.88 ± 0.08** | **0.65 ± 0.09** |
| Out-of-distribution unseen dataset | | | | |
| UNETR | 0.73 ± 0.07 | 7.86 ± 5.40 | **0.85 ± 0.08** | 0.68 ± 0.16 |
| MAPL3 | **0.77 ± 0.04** | **5.32 ± 2.60** | 0.82 ± 0.12 | **0.75 ± 0.14** |

**Supplementary Table 2.** SPECTRE framework ablation study

| Experiment | DSC | HD95 | Recall | Precision |
|---|---|---|---|---|
| No preprocessing | 0.69 ± 0.10 | 5.77 ± 2.55 | 0.63 ± 0.20 | **0.86 ± 0.17** |
| Voxel-wise adaptive bias | 0.68 ± 0.11 | 6.13 ± 3.36 | 0.67 ± 0.21 | 0.84 ± 0.20 |
| Patch- and voxel-wise adaptive bias | 0.72 ± 0.10 | 5.90 ± 3.54 | 0.72 ± 0.19 | 0.82 ± 0.18 |
| No MIP and edge-weighted loss | 0.71 ± 0.12 | 7.42 ± 4.39 | 0.77 ± 0.16 | 0.75 ± 0.21 |
| No edge-weighted loss | 0.73 ± 0.11 | 6.15 ± 3.89 | 0.76 ± 0.18 | 0.79 ± 0.21 |
| MAPL3 | **0.77 ± 0.09** | **4.20 ± 3.09** | **0.81 ± 0.13** | 0.79 ± 0.17 |

**Supplementary Table 3.** The effects of self-supervised pretraining on SPECTRE performance.

| Algorithm | DSC | HD95 | Recall | Precision |
|---|---|---|---|---|
| In-distribution test dataset | | | | |
| Scratch | 0.74 ± 0.04 | 4.27 ± 1.73 | **0.88 ± 0.08** | 0.65 ± 0.09 |
| Pre-trained | **0.77 ± 0.02** | **3.86 ± 1.64** | 0.83 ± 0.10 | **0.73 ± 0.07** |
| Out-of-distribution unseen dataset | | | | |
| Scratch | 0.77 ± 0.04 | 5.32 ± 2.60 | **0.82 ± 0.12** | 0.75 ± 0.14 |
| Pre-trained | **0.77 ± 0.06** | **5.28 ± 3.01** | 0.75 ± 0.16 | **0.82 ± 0.13** |

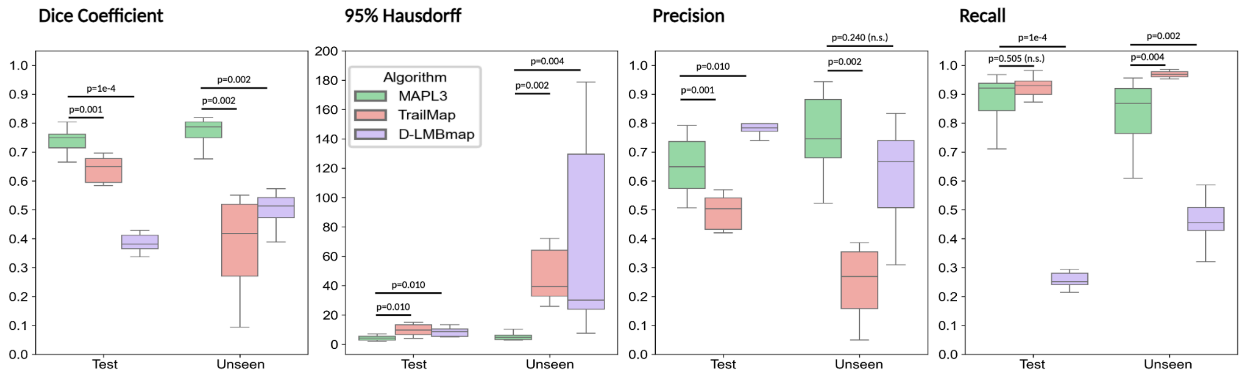


**Supplementary Data Fig. 1.** Performance of SPECTRE trained from random initialization compared with fine-tuned benchmark models. SPECTRE without self-supervised pretraining (MAPL3; green), TrailMap (red), and D-LMBmap (purple) were trained using the same annotated training dataset and evaluated on the test and unseen datasets. Segmentation performance was assessed using the Dice similarity coefficient, 95th-percentile Hausdorff distance (HD95), precision, and recall. SPECTRE achieved significantly higher Dice similarity coefficients and lower HD95 values than both fine-tuned benchmarks across the test and unseen datasets. TrailMap achieved recall comparable to or higher than SPECTRE but exhibited substantially lower precision, whereas D-LMBmap achieved comparable precision on the unseen dataset but substantially lower recall. Box plots show the median, interquartile range, and minimum and maximum values. Exact P values are shown above the corresponding comparisons; n.s., not significant.

.

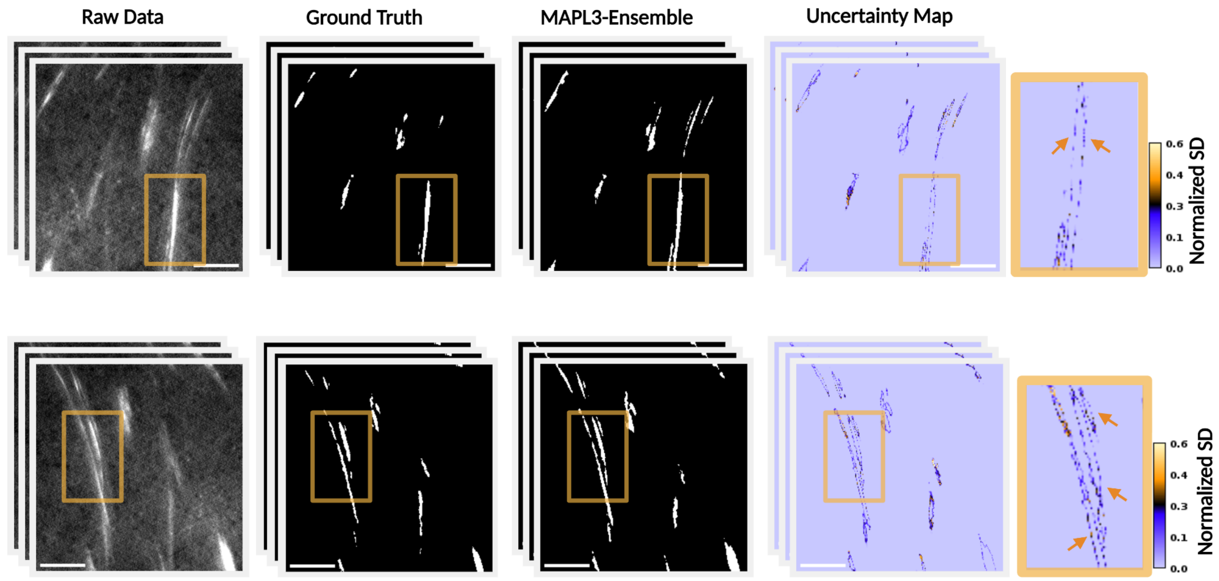

**Supplementary Data Fig. 2.** Monte Carlo dropout provides spatially resolved estimates of segmentation uncertainty. Representative LSFM patches showing the raw fluorescence signal, expert-corrected reference annotation, MAPL3 MC-dropout ensemble segmentation, and corresponding voxel-wise uncertainty map. The ensemble segmentation was generated from the mean predicted probability across 50 stochastic forward passes with dropout retained during inference. Predictive uncertainty was calculated as the voxel-wise standard deviation across the 50 probability maps and normalized for visualization. Enlarged views of the boxed regions show elevated uncertainty along axonal boundaries and at locations where the predicted axonal extent varied across stochastic passes, indicated by orange arrows. Scale bars are shown in white. The uncertainty maps represent epistemic variability in SPECTRE predictions and do not capture all sources of uncertainty arising from image noise, fluorescence variability, or reference-annotation ambiguity.

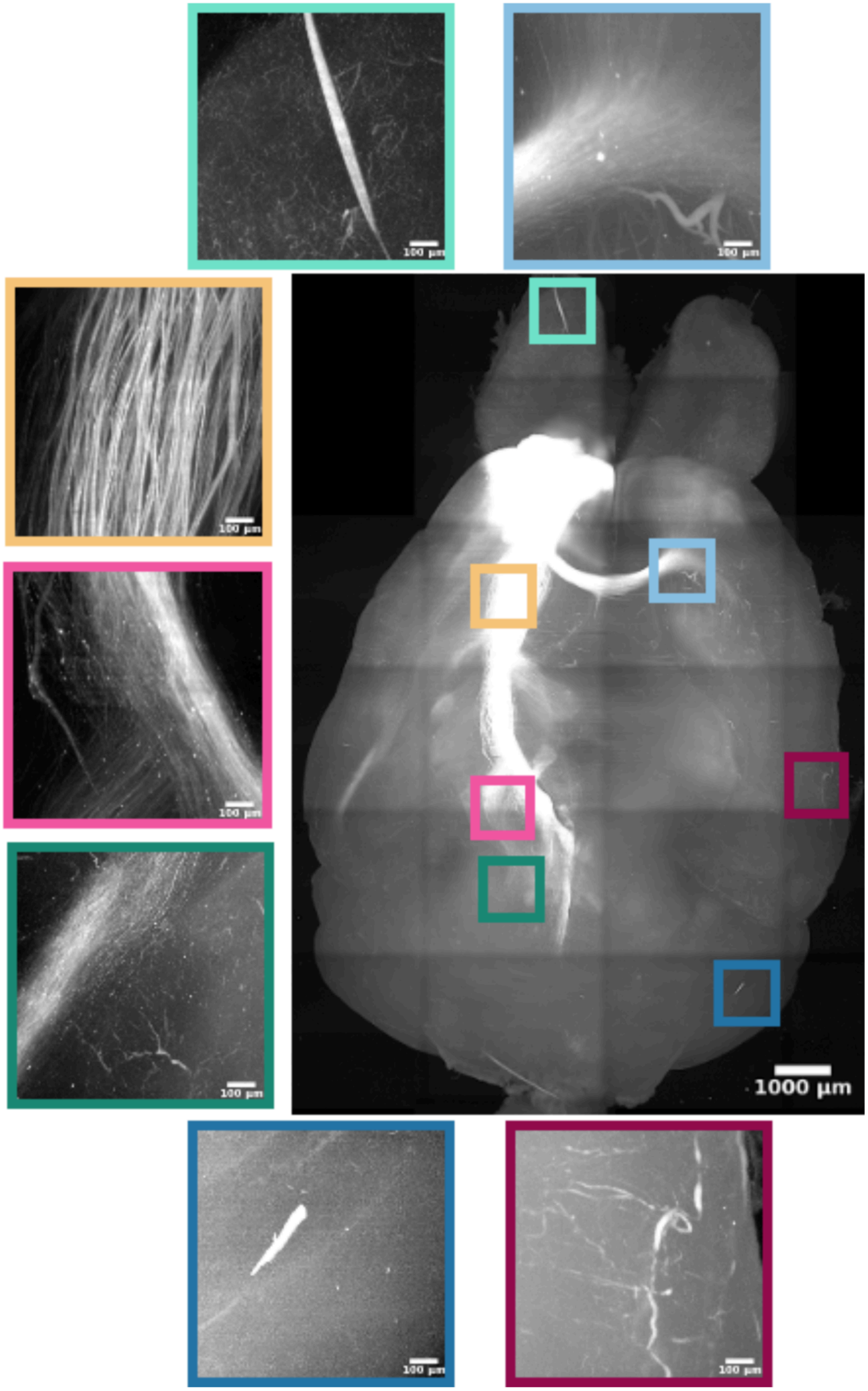


**Supplementary Data Fig. 3.** Whole-brain maximum intensity projection of axonal fibers originating from the right orbitofrontal cortex. Selected 3D sub-volumes ($512^3$ voxels) are shown to illustrate regions with varying axonal densities, imaging artifacts, and mixed signal profiles, highlighting the challenges of accurately mapping these fibers across the whole brain.

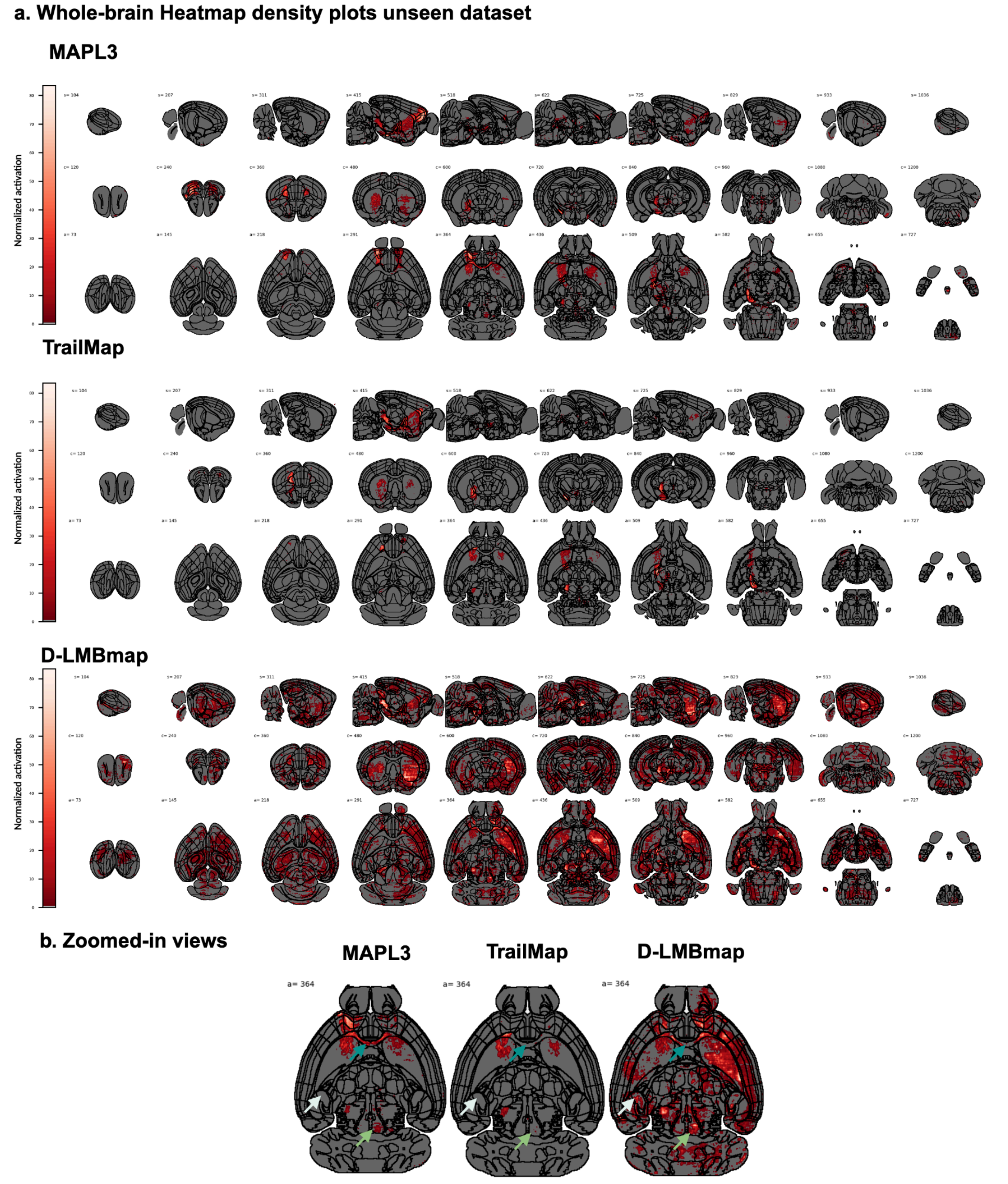


**Supplementary Data Fig. 4. a.** Whole-brain axonal projections from the right orbitofrontal cortex of a representative subject in the high-resolution unseen dataset (1.8x1.8x3 $\mu m^3$ isotropic scanned on a SmartSPIM LSFM), reconstructed by MAPL3, TrailMap, and D-LMBmap. Voxelized segmentation maps were warped to the Allen atlas (10 µm), and region boundaries were overlaid for comparison. Rows 1–3 show sagittal, axial, and coronal views at randomly chosen cut planes. **b.** Enlarged views of selected regions. Dark green arrow: corpus callosum, accurately reconstructed by MAPL3 but missed by TrailMap and fragmented in D-LMBmap. Light green arrow: periaqueductal gray (PAG), where MAPL3 detected localized fibers that were missed by TrailMap and oversegmented by D-LMBmap. White arrow: basolateral amygdalar nucleus, which does not receive orbitofrontal input and was correctly identified as lacking projections by MAPL3 and TrailMap, but falsely reconstructed by D-LMBmap

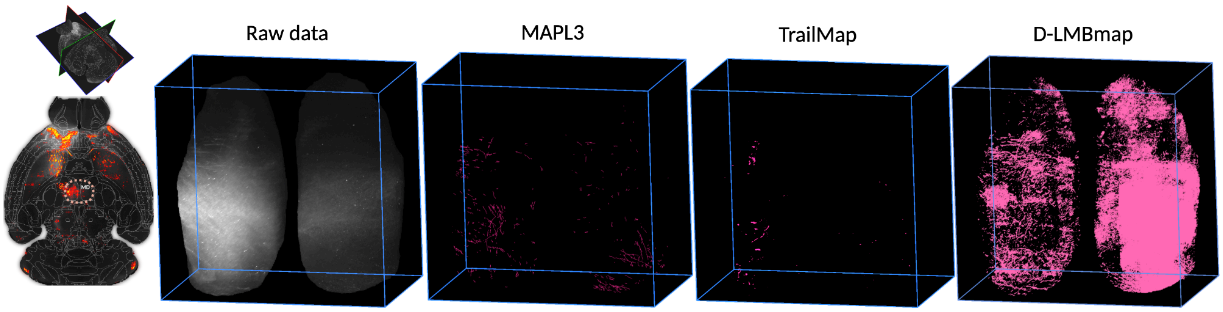


**Supplementary Data Fig. 5. Maximum-intensity rendering of axonal fibers in the mediodorsal thalamic nucleus (MD) in native space.** Allen labels were warped to the native space of a randomly selected subject from the application dataset using the deformation matrix obtained by the registration module. Raw data and segmentation maps generated by MAPL3, TrailMap, and D-LMBmap are shown. MAPL3 accurately detected sparse, localized axonal fibers in the MD, which were missed by TrailMap and oversegmented by D-LMBmap.

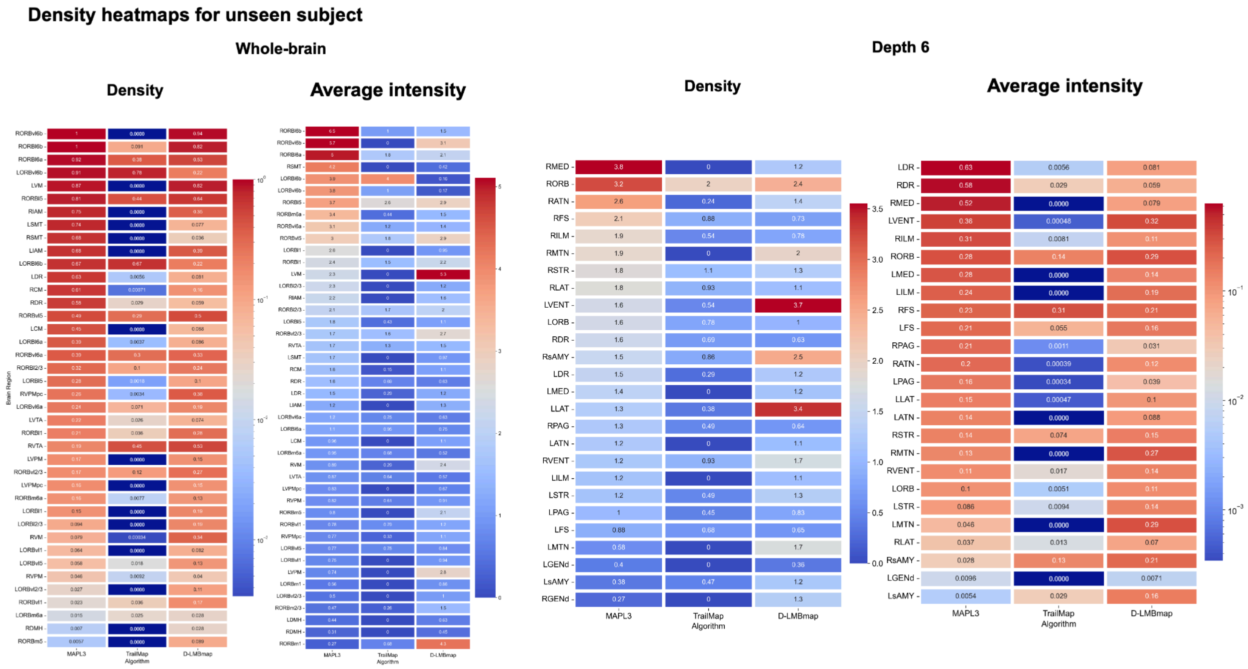


**Supplementary Data Fig. 6.** Whole-brain heatmaps of axonal density and voxel-wise average intensity of the warped segmentation map originating from the seed region (right orbitofrontal cortex), reconstructed by MAPL3, TrailMap, and D-LMBmap, for an example low-resolution unseen dataset (5µm isotropic, scanned on a LaVision LSFM). Projections are shown from key regions and their subdivisions (whole-brain analysis) and depth 6, including the orbitofrontal cortex (ORB), striatum, ventral tegmental area (VTA), dorsal thalamus, hypothalamic nuclei, and periaqueductal gray (PAG).

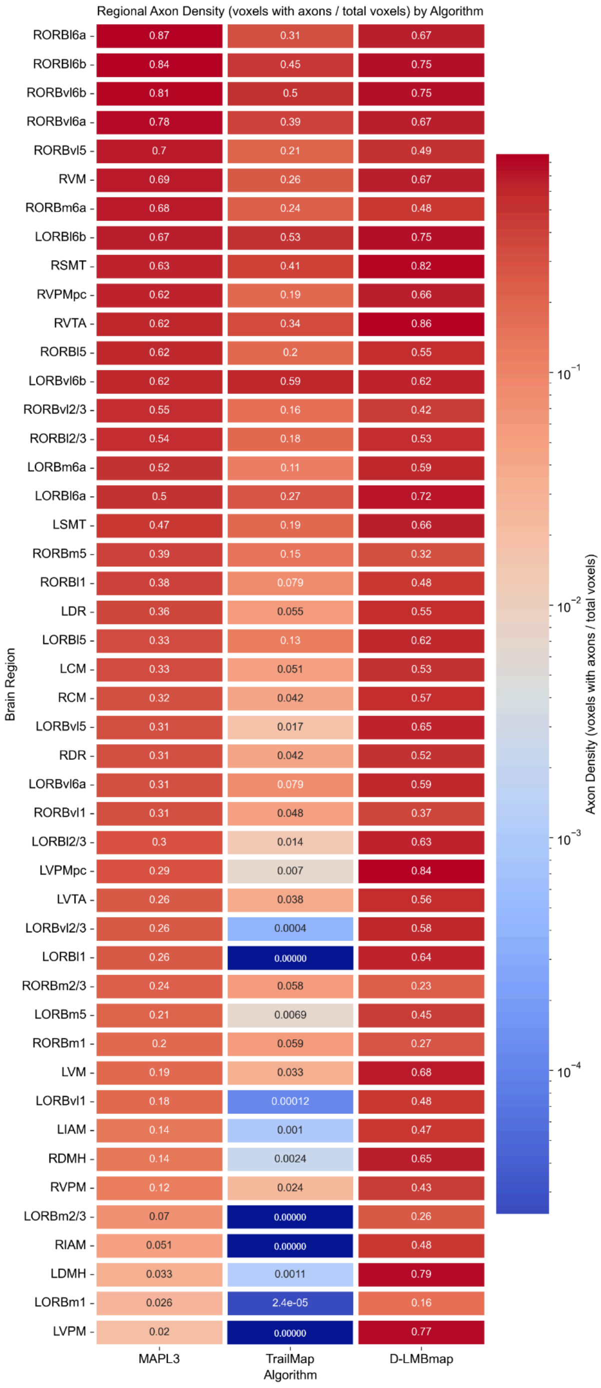


**Supplementary Data Fig. 7.** Whole-brain heatmaps of axonal density of the high-resolution unseen dataset (1.8x1.8x3 μm$^3$ isotropic scanned on a SmartSPIM LSFM) originating from the seed region (right orbitofrontal cortex), reconstructed by MAPL3, TrailMap, and D-LMBmap, averaged across all subjects. Projections are shown from key regions and their subdivisions, including the orbitofrontal cortex (ORB), striatum, ventral tegmental area (VTA), dorsal thalamus, hypothalamic nuclei, and periaqueductal gray (PAG).

**a. Key-regions with sub-layers density comparison**

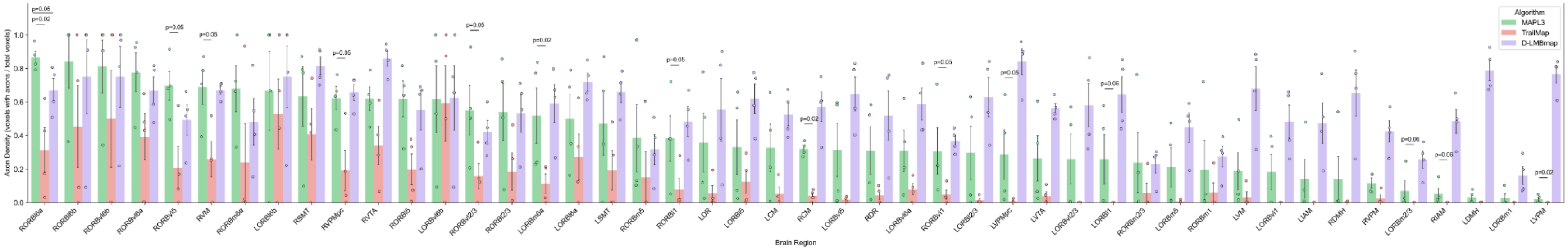


**b. Key-regions with sub-layers average intensity of voxelized and warped segmentation maps comparison**

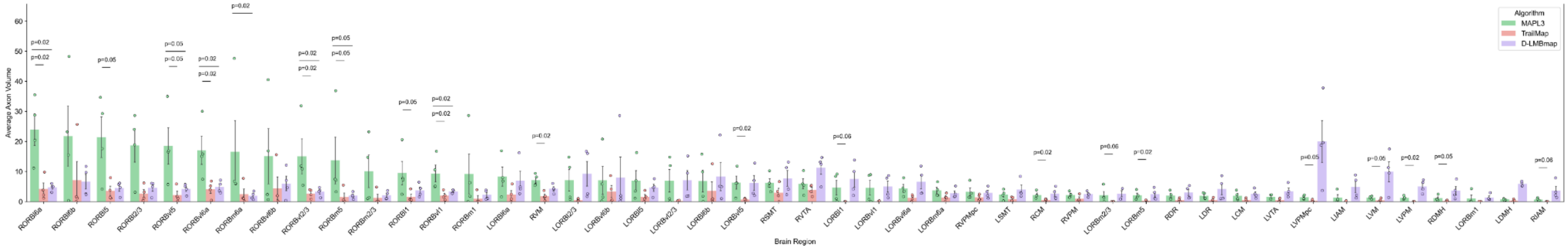


**Supplementary Data Fig. 8.** Whole-brain axonal density maps (a) and bar plots of voxel-wise average intensity of the warped segmentation map (b) of the high-resolution unseen dataset (1.8x1.8x3 $\mu m^3$ isotropic scanned on a SmartSPIM LSFM) from seed projections originating in the right orbitofrontal cortex, reconstructed with MAPL3, TrailMap, and D-LMBmap. Projections are quantified across major target regions and their subdivisions, including the orbitofrontal cortex (ORB), striatum, ventral tegmental area (VTA), dorsal thalamus, hypothalamic nuclei, and periaqueductal gray (PAG). Bar plots show mean values ± standard error across subjects; statistical significance was assessed using a two-sided Mann–Whitney U test.

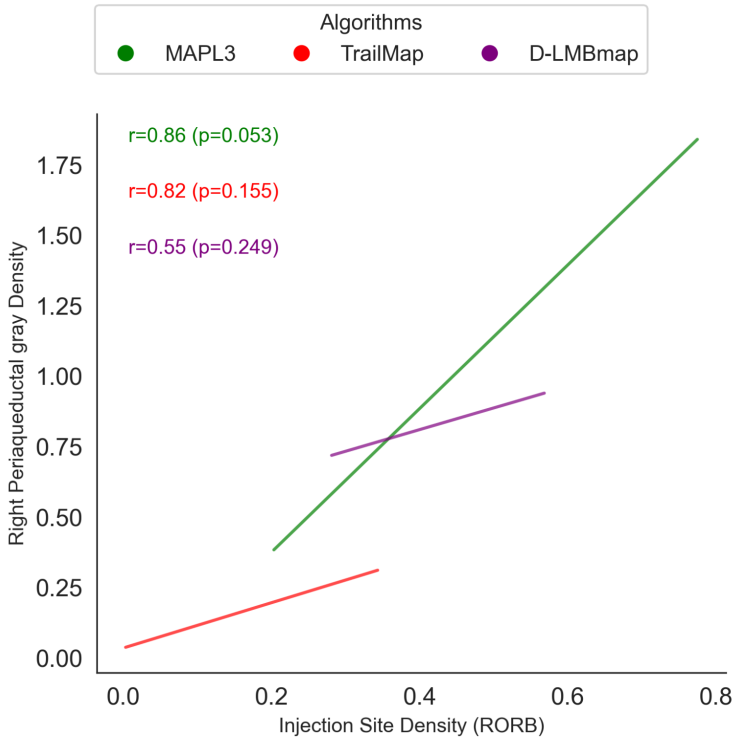


**Supplementary Data Fig. 9.** Correlation between injection-site density and right PAG fiber density at depth 6 (based on bootstrapping, n=1000 permutations). MAPL3 demonstrates a positive relationship (r = 0.86, p = 0.053), whereas TrailMap and D-LMBmap fail to capture this trend.

**Whole-brain rendering of streamlines**

Raw data downsampled and warped to Allen

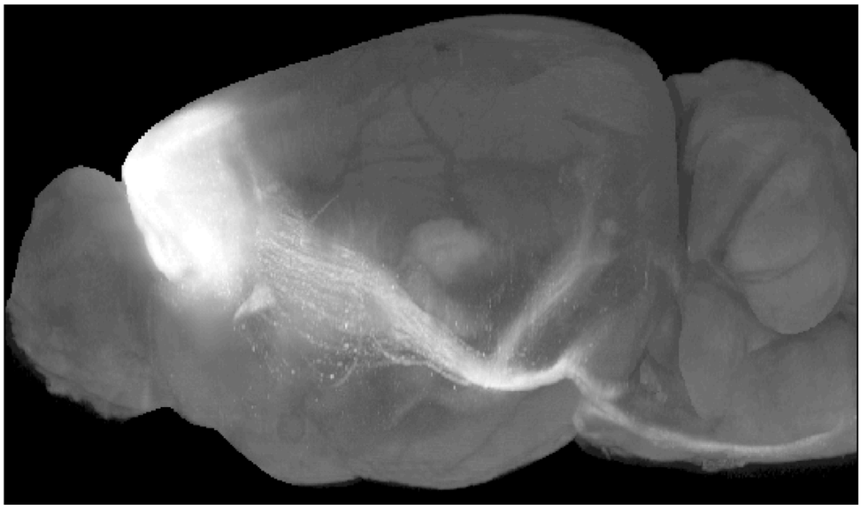

MAPL3

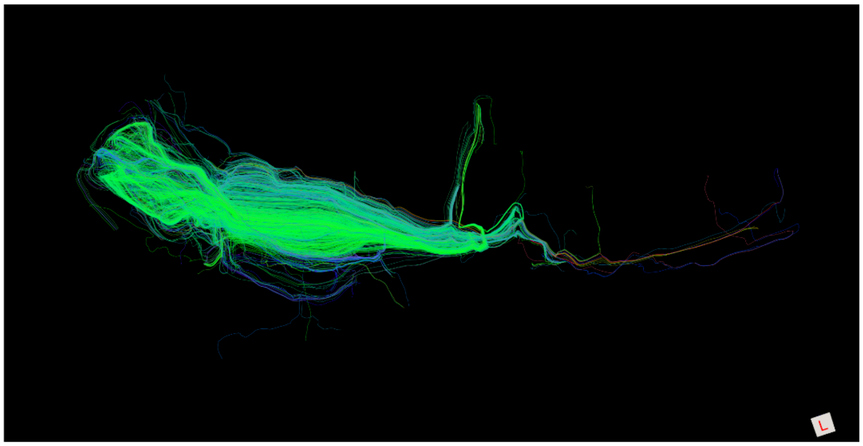

STA on raw data

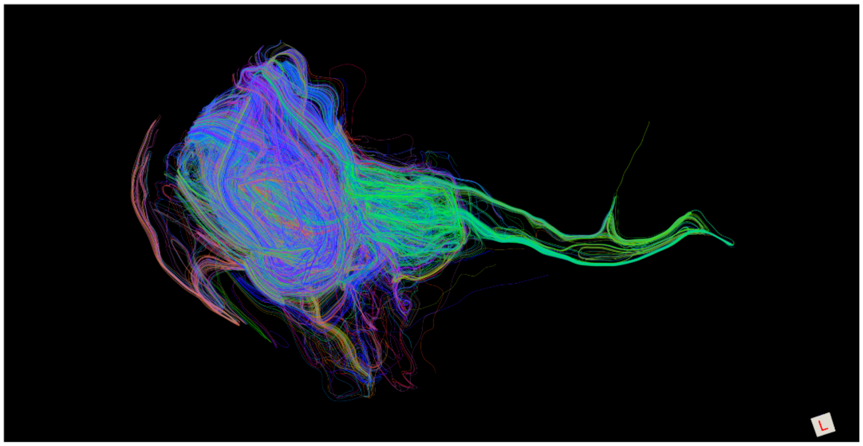

**Supplementary Data Fig. 10.** Whole-brain visualization of RORB-seeded streamlines reconstructed from MAPL3 segmentation and downsampled fluorescence data. Top, sagittal projection of the downsampled LSFM fluorescence signal registered to the Allen Reference Atlas. Middle and bottom, whole-brain renderings of streamlines seeded from the RORB and reconstructed using the MAPL3 tractography module or applying STA analysis directly to the downsampled and atlas-registered LSFM fluorescence signal, respectively. MAPL3 segmentation-based tractography reconstructed a more extensive set of putative projection trajectories extending from the injection site toward distal cortical, subcortical, and midbrain regions, whereas tractography performed directly on the downsampled fluorescence signal produced a more restricted tractogram. Streamline colors facilitate visualization of individual reconstructed trajectories. Streamlines represent computational estimates derived from local orientation fields and should not be interpreted as one-to-one reconstructions of individual biological axons.

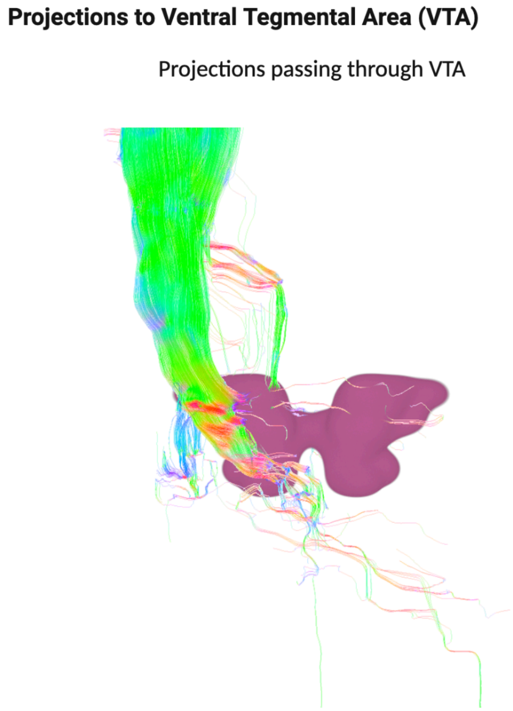


**Supplementary Data Fig. 11**. Streamline-wise separation of trajectories traversing and ending within the ventral tegmental area. Three-dimensional visualization of RORB-seeded streamlines annotated according to their relationship with the ventral tegmental area (VTA; magenta). Left, streamlines passing the VTA and continuing toward other brain regions. Right, streamlines with distal reconstructed endpoints located within the VTA. This region-specific filtering illustrates how MAPL3 distinguishes regional streamline traversal from distal endpoint localization. Streamline endpoints represent computationally reconstructed locations and should not be interpreted as direct evidence of biological axon terminals or synaptic connectivity.

### Quantification of streamlines in key-regions with sub-layers comparison

#### Tract density

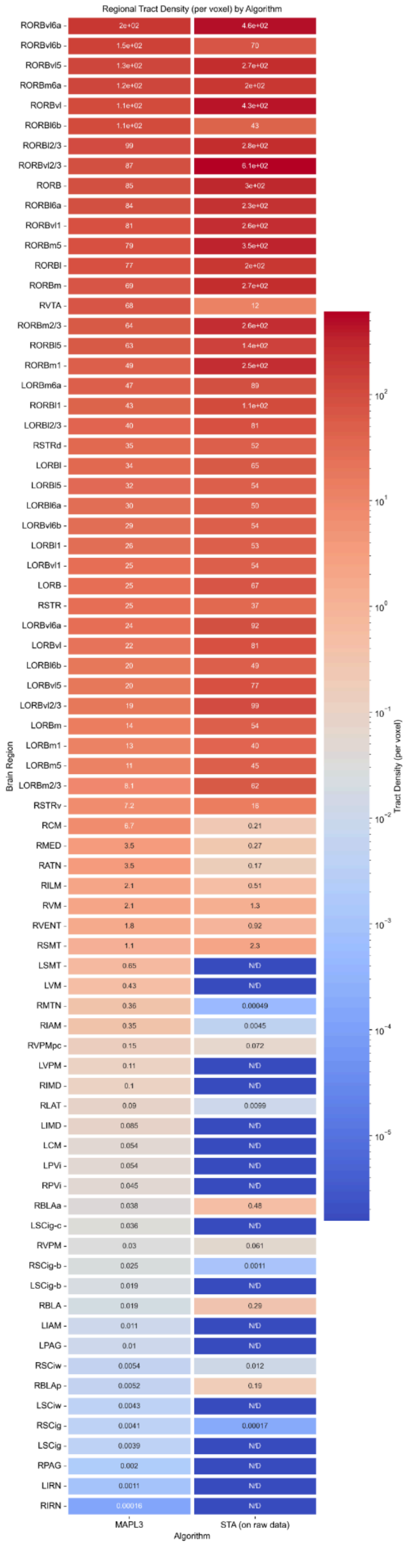


#### Endpoint density

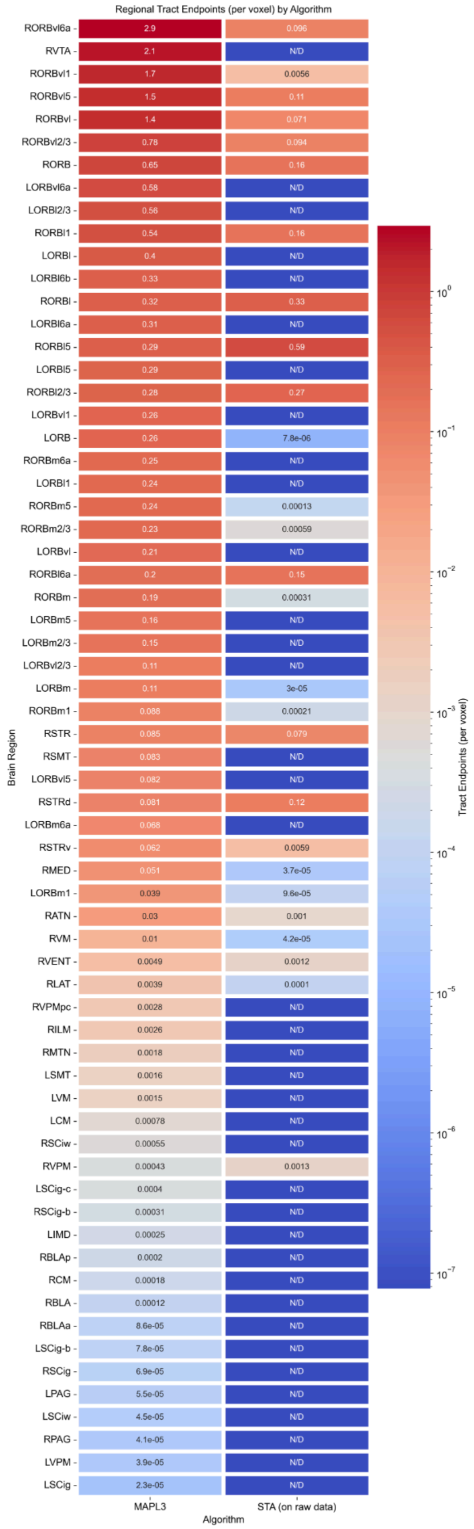

**Supplementary Data Fig. 12.** Regional quantification of RORB-seeded streamlines reconstructed using MAPL3 segmentation-based and raw-signal–based tractography. Heatmaps compare streamline tract density (left) and distal streamline endpoint density (right) across atlas-defined brain regions and subregions in the high-resolution application dataset. Streamlines were seeded from the right orbitofrontal cortex (RORB) and reconstructed using either structure tensor analysis applied to the voxelized MAPL3 segmentation map or the same analysis applied directly to the downsampled LSFM fluorescence signal registered to Allen Reference Atlas space. Values represent the number of streamline traversals or distal endpoints per voxel within each region and are displayed on logarithmic color scales. MAPL3 segmentation-based tractography produced more extensive regional tract and endpoint profiles, particularly in distal regions where raw-signal–based tractography yielded no detectable streamlines or endpoints. N/D, not detected; ARA, Allen Reference Atlas; STA, structure tensor analysis.

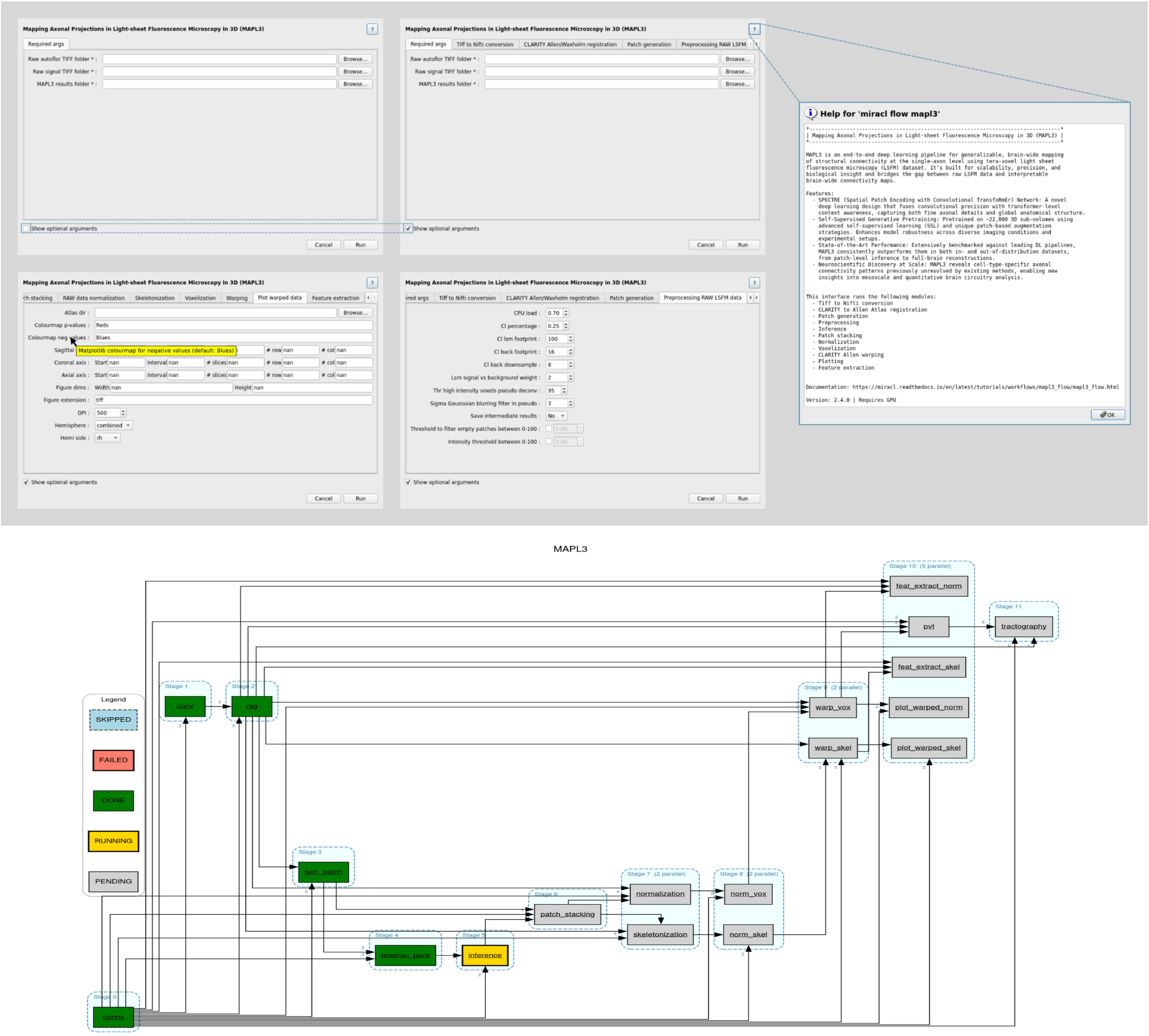


**Supplementary Data Fig. 13.** MAPL3's graphical user interface (GUI; Top) and a representative output directed acyclic graph (DAG; bottom) of the different modules for an example run.
